# Inhibition of glioblastoma cell proliferation and invasion by the choline-kinase inhibitor JAS239 varies with cell type and hypoxia

**DOI:** 10.1101/2024.01.17.576078

**Authors:** Claire Louise Kelly, Martyna Wydrzynska, Marie M Phelan, Sofya Osharovich, Edward J. Delikatny, Violaine Sée, Harish Poptani

**Author notes:** **Corresponding and co-senior authors Corresponding authors Email:**. Present address: University Claude Bernard Lyon1, CNRS UMR 5305, Tissue Biology and Therapeutic Engineering Laboratory (LBTI), 69007 Lyon, France. Shared last authorship (Violaine Sée, Harish Poptani).

## Abstract

**Background:** Elevated choline kinase alpha (ChoK) is observed in most solid tumours including glioblastomas (GBM), yet until recently, inhibitors of ChoK have demonstrated limited efficacy in GBM models. Given that hypoxia is associated with GBM therapy resistance, we hypothesised that tumour hypoxia could be responsible for such limitations. We therefore evaluated in GBM cells, the effect of hypoxia on the function of JAS239, a potent ChoK inhibitor.

**Methods:** Rodent (F98 and 9L) and human (U-87 MG and U-251 MG) GBM cell lines were subjected to 72 hours of hypoxia conditioning and treated with JAS239 for 24 hours. NMR metabolomic measurements and analyses were performed to evaluate the signalling pathways involved. In addition, cell proliferation, cell cycle progression and cell invasion were measured in cell monolayers and 3D spheroids, with or without JAS239 treatment in normoxic or hypoxic cells to assess how hypoxia affects JAS239 function.

**Results:** Hypoxia and JAS239 treatment led to significant changes in the cellular metabolic pathways, specifically the phospholipid and glycolytic pathways associated with a reduction in cell proliferation via induced cell cycle arrest. Interestingly, JAS239 also impaired GBM invasion. However, JAS239 effects were variable depending on the cell line, reflecting the inherent heterogeneity observed in GBMs.

**Conclusion:** Our findings indicate that JAS239 and hypoxia can deregulate cellular metabolism, inhibit proliferation and alter cell invasion. These results may be useful for the design of new therapeutic strategies based on ChoK inhibition that can act on multiple pro-tumorigenic features.

## Introduction

Primary glioblastoma (GBM) are the most common adult primary brain tumours with current standard of care being ineffective at curing the patient in most cases.[1, 2] Survival ranges between 14–16 months after diagnosis with a 5-year survival rate of less than 5%, making GBM the deadliest brain tumour in adults.[3–5] Currently temozolomide, an alkylating agent, and bevacizumab, a vascular endothelial growth factor receptor inhibitor, are the only food and drug administration of USA (FDA) approved systemic therapies for newly diagnosed and recurrent GBM, respectively.[6–8] Targeted therapy for GBM has yet to demonstrate a substantial survival benefit, therefore there is an urgent need to develop novel therapies for these patients.

Molecular phenotypes of distinct regions within the same tumour can be diverse and generate unique tumour microenvironments.[9–12] For example, a low oxygen (hypoxic) environment initiates specific genetic and molecular alterations, mediated via the hypoxia inducible transcription factor (HIF).[9, 13] This cellular adaptation response contributes to enhanced cell survival, resistance to therapies and intratumoural heterogeneity.[14] The specific impact of hypoxia on the outcome of drug efficacy, combined with intratumoural heterogeneity, generates a variable tumour sensitivity to targeted therapies. Metabolic inhibitors can reduce therapy resistance by limiting the levels of critical metabolites necessary for DNA damage repair, enhancing chemo- and radiotherapy sensitivity.[15], thereby supporting the rationale for their use as potential adjuvants to chemotherapy.

Choline kinase (ChoK) is the first committed step of the Kennedy pathway, for phosphatidylcholine synthesis, the most abundant phospholipid in eukaryotic cell membranes.[16, 17] Inhibition of ChoKα has shown promising results in *in vivo* models of breast cancer and more recently in GBM.[18–20] However, given that hypoxia is associated with GBM therapy resistance, we hypothesised that tumour hypoxia could be a contributing factor for the limited efficacy observed to date. We therefore evaluated the effect of hypoxia on the function of JAS239, a potent ChoK inhibitor, in four of the most commonly used GBM cell lines.

Moreover, when assessing the effects of hypoxia on drug efficiency *in vitro*, most studies use a standard 24h pre-incubation hypoxic conditions, which is too short to recapitulate the long-term cellular adaptation that occurs within hypoxic tumours.

We here investigated the metabolic changes and the effects on cell proliferation and invasion triggered by JAS239 as well as the impact of prolonged hypoxia (up to 96 hours) on these mechanisms. We used 2D cell culture to investigate the JAS239 effects on cell proliferation, and 3D spheroid models to assess the effects on cell invasion, using four different GBM cell lines from rat and human origin.

## Methods

### Cell lines

All GBM cell lines (F98, 9L, U-87 MG and U-251 MG) were maintained at 37°C in a 5% CO₂ humidified atmosphere (see **Supplementary Section 1** for details). Cells were passaged every 2‒3 days at 80‒90% confluency up to passage 10. For hypoxic preconditioning, cells were incubated for 72 hours (h) at 1% O_2_ (hypoxia) in a hypoxic workstation (Don Whitley Hypoxic Workstation, England), prior to drug treatment. Once treated, cells were incubated for a further 24 h during drug exposure. Drug treatment was performed within the workstation without reoxygenation of the cells.

### Stable Cell lines

F98 and U-87 MG histone H2B monomeric red fluorescent protein (H2BmRFP) and F98 and 9L Luc Zs Green stable cell lines were generated for 3D spheroid experiments to enable cell tracking (see **Supplementary Section 1** for full experimental details).

### Cell Viability

Cell viability was assessed using the 3-(4,5-Dimethylthiazol-2-yl)-2,5-diphenyltetrazolium bromide (MTT) assay (see **Supplementary Section 1** for full experimental details). Background absorbance was subtracted from all conditions and then plotted as a percentage of DMSO control to determine the IC50 for each cell line.

### NMR on cell extracts

Aqueous metabolites were extracted from cell samples and 1D ^1^H NMR spectra were acquired (See **Supplementary Section 1** for full experimental details). Multivariate analysis was performed, including principal component analysis (PCA) and partial least square discriminant analysis (PLS-DA). To elucidate the relevance of metabolites and metabolic pathways, MetaboAnalyst 5.0 was used for pathway enrichment analysis.[21]

### Western Blotting

20–30 µg protein were resolved on a 10% SDS-polyacrylamide gel, transferred onto a nitrocellulose membrane and probed with primary antibodies (anti-choline kinase alpha [Abcam; ab88053], anti-vinculin [Abcam; ab129002], anti-tubulin [Cell Signalling; 2128S], anti- E2F5 [Santa Cruz; sc-1083], anti-E2F1 [Cell Signalling; 3742], anti- α3+β1 [Abcam; ab217145], anti-HIF-1α [Protein Tech; 20960-1-AP], anti-HIF-2α [Bethyl Labs; A700-003]) at 4°C for 12 h. Membranes were incubated with anti-rabbit (1:10,000) horseradish peroxidase-linked secondary antibody (Cell Signalling) for 2 h at room temperature. Signal was developed using Amersham ECL Prime Western blotting Detection reagent (GE Healthcare, Chicago, IL, USA), and images taken using a G:BOX gel imaging system (Syngene, Cambridge, UK).

### Flow Cytometry

Cells were washed with phosphate-buffered saline (PBS), trypsinised and resuspended in PBS at 0.5–1x10^6^/500 µl. Cells were fixed by adding them dropwise to ice-cold 70% ethanol whilst gently vortexing, and incubated for 30 min at 4°C. Samples were washed 2x in ice-cold PBS and centrifuged for 10 min at 300 xg. 1 ml of 0.5% Tween20/PBS was added to sample pellets and incubated at room temperature for 5 min. Samples were washed 2x with ice-cold PBS, centrifuged for 3 min at 300 xg and all supernatant removed. Immediately prior to data acquisition, samples were re-suspended in 1 ml 1 µg/ml Hoechst 33342 in PBS (Invitrogen 10 mg/ml stock). Cell cycle analysis was performed using a BD Bioscience FACSCanto II flow cytometry system using a 401 nm laser. Flow cytometry data was analysed using NovoExpress 1.4.1 software. The population of cells analysed were gated according to **Supplementary** Figure 1 to ensure only live singlets were included in the analysis and NovoExpress built-in cell cycle analysis software was used to quantify the proportion of cells in G0/G1, S and G2/M phase.

### 3D spheroids

1x10^6^ F98 H2B RFP or U-87 MG H2B RFP cells were seeded in 1ml filtered culture media in one well of a 5D spherical plate (Kugelmeiers, Switzerland) and incubated at 37°C overnight for spheroid formation. Spheroids were imaged on a Zeiss Light-Sheet microscope and analyzed using using Imaris v9.6 software (www.imaris.oxinst.com; Oxford Instruments, UK). See **Supplementary Section 1** for full experimental details.

## Results

### JAS239 efficacy varies by cell line in response to hypoxic conditioning

Since hypoxia influences drug effectiveness in various cancer models,[22–24] we firstly assessed the effects of JAS239 on GBM cell metabolic activity under 96 h of hypoxic exposure (72 h of hypoxic preconditioning followed by 24 h JAS239 treatment under hypoxic conditions). No significant difference in IC50 was found between normoxic (21% O_2_) and hypoxic (1% O_2_) conditioning across all four GBM cell lines upon 24 h of JAS239 treatment (**Supplementary** Figure 2). Similar results were observed in cells preconditioned for 72 h but then reoxygenated for 24 h during JAS239 treatment (red lines). This experiment was designed to check if there was a direct chemical effect of the lack of oxygen on drug efficacy as it has been reported for other drugs such as phleomycin.[25] JAS239 remained active and effective under all oxygenation patterns. JAS239 was also shown to increase *CHK* gene expression and ChoKα protein expression in F98 and 9L normoxic cells only (**Supplementary** Figure 3 **A, B & C**).

Conversely, *CHK* gene expression increased in U-87 MG and U-251 MG normoxic and hypoxic cells while ChoKα protein expression decreased with JAS239 treatment (**Supplementary** Figure 3 **A, D & E**). Hypoxic exposure was monitored by measuring HIF-1α protein levels. HIF-1α was indeed stabilised within 8 h of hypoxic exposure and was maintained throughout 72 h in F98, 9L and U-87 MG cells indicating that cells were responding well to hypoxia (**Supplementary** Figure 4 **A & B**). Surprisingly, U-251 MG cells showed HIF-1α stabilisation in normoxic conditions as well as hypoxic conditions. HIF-2α protein levels were also increased in the four cell lines in hypoxia (**Supplementary** Figure 4 **B & D**). For both rat cell lines (F98 and 9L), there was detectable HIF-2α protein in normoxic conditions. Interestingly treatment with JAS239, after 72 h of hypoxia, reduced HIF-1α and HIF-2α levels in all cell lines.

### Metabolite enrichment in JAS239 normoxic and hypoxic cells

We further established the intracellular metabolite modifications induced by JAS239 treatment. We focused on GBM metabolism, using NMR-based metabolomic analysis since JAS239 targets the choline pathway by competitively binding to the active site of ChoKα, preventing phosphorylation of choline during phosphatidylcholine biosynthesis.[16, 18] We therefore hypothesised that phosphocholine levels would be reduced in JAS239 treated cells. The effect of prolonged hypoxia on phosphocholine levels was unknown. Upon 6 h of JAS239 treatment, a clear reduction in phosphocholine and an increase in glycerophosphocholine was detected in both normoxic and hypoxic conditions, as shown on the representative spectra of F98 cells (**Figure 1 A**). Data transformation using PCA analysis exhibited variance due to oxygenation as opposed to JAS239 treatment for all but U-251 MG cell line and the loadings of metabolites that were most influential in the first and second principal components were identified and shortlisted (representative F98 PCA loadings are shown in **Figure 1 B**). PCA analysis of the F98 (**Figure 1 B**) and 9L (**Supplementary** Figure 5 **A**) cells showed that principal components 1 and 2 overlapped. Phosphocholine, formate, and glycerophosphocholine ranked most influential in PCA loadings of F98 cells, whilst acetate, phosphocholine and creatine ranked most influential in PCA loadings of 9L cells, however, this was most likely due to hypoxia alone rather than JAS239 treatment. Similarly, PCA analysis of U-87 MG cells (**Supplementary** Figure 6 **A**) demonstrated separation between principal components 1 and 2 due to oxygenation, with glycerophosphocholine, formate and choline ranking most influential, while the U-251 MG PCA analysis of principal components 1 and 2 completely overlapped (**Supplementary** Figure 7 **A**). The shortlisted metabolites were then used to perform a PLS-DA analysis between treatment and hypoxic conditions (representative F98 **Figure 1 C**). The PLS-DA models were optimally fit with four/five components and yielded well-defined separation between JAS239 and DMSO as well as hypoxic and normoxic conditions with component 5 ROC scores ranging for each group between 0.91–1 (representative F98 **Figure 1 C**). F98 and 9L cells demonstrated clear discrimination between hypoxia/normoxia and DMSO/JAS239 treatment in both cell lines. As hypothesised, phosphocholine glycerophosphocholine and choline were listed amongst the VIP loading plots for F98 cells (**Figure 1 D**) and phosphocholine and choline for 9L cells but with differing influence, nonetheless indicating this pathway is influential in the separation of treatment conditions. Although the PLS-DA plot of components 1 and 2 of the 5-component models for both U-87 MG and U-251 MG data sets (**Supplementary** Figures 5 **B** & **7 B**) do not show visible discrimination between groups, discrimination was visible in higher components of the model (U-87 MG component 5 ROCs: 0.74–1 and U-251 MG component 5 ROCs: 0.97–1).

**Figure 1.**
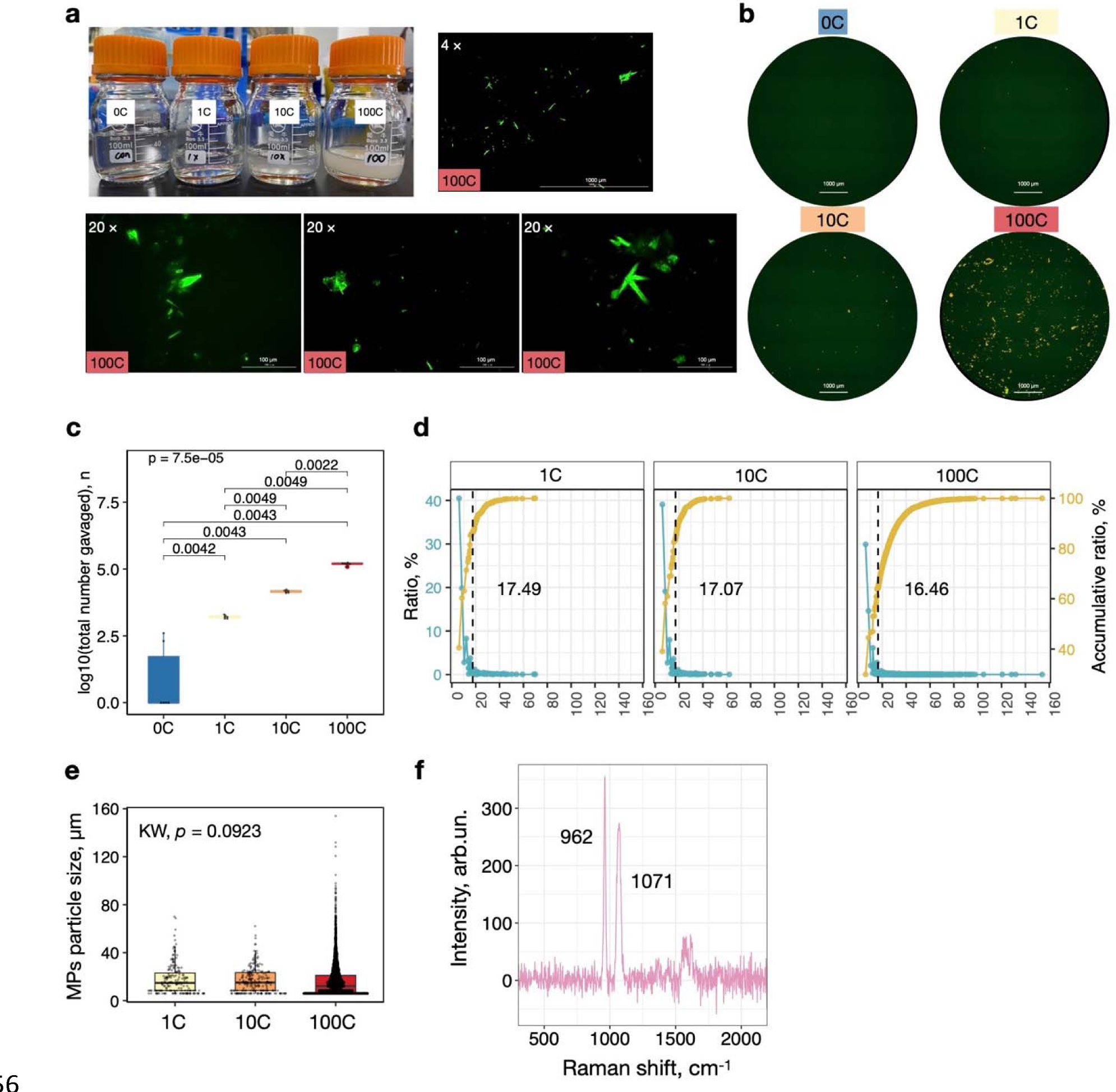
Representative figures for the metabolomic analysis pipeline. A) Example F98 ^1^H 1D NMR spectra acquired at 700MHz and referenced to internal standard (TSP). B) PCA analysis. C) PLS-DA analysis (component 5 ROC scores for each group vs all other groups: H_D = 1; H_J = 1; N_D = 0.91 and N_J = 1) and D) most influential metabolites. Metabolites with VIP >2 (D) informed the metabolite set enrichment analysis of top 20 metabolites using the hypergeometric test E). D, DMSO; H, hypoxia; J; JAS239; N, normoxia; PCA, principal component analysis; PLS-DA, partial least square discriminant analysis; ROC, receiver operating characteristic; VIP, variable importance in projection coefficients.

From the PLS-DA analysis, a shortlist of metabolites with a VIP score >1 was used to perform metabolite enrichment analysis (representative F98 **Figure 1 E**) and the top 5 most influential metabolomic pathways altered in response to JAS239 and hypoxia are shown in **Figure 2** for all cell lines. Enrichment analysis of F98 cells highlights the influential metabolites associated mostly with lipid metabolism pathways, particularly phospholipid metabolism. Conversely, enrichment analysis of metabolites most influential in 9L cells ranked the ‘Warburg effect’ followed by mitochondrial metabolism or the tricarboxylic acid (TCA) cycle. Pathway enrichment of U-87 MG data ranked the phosphatidylethanolamine metabolism as most influential, clearly indicating cell line specificity in the response to JAS239 treatment.

**Figure 2.**
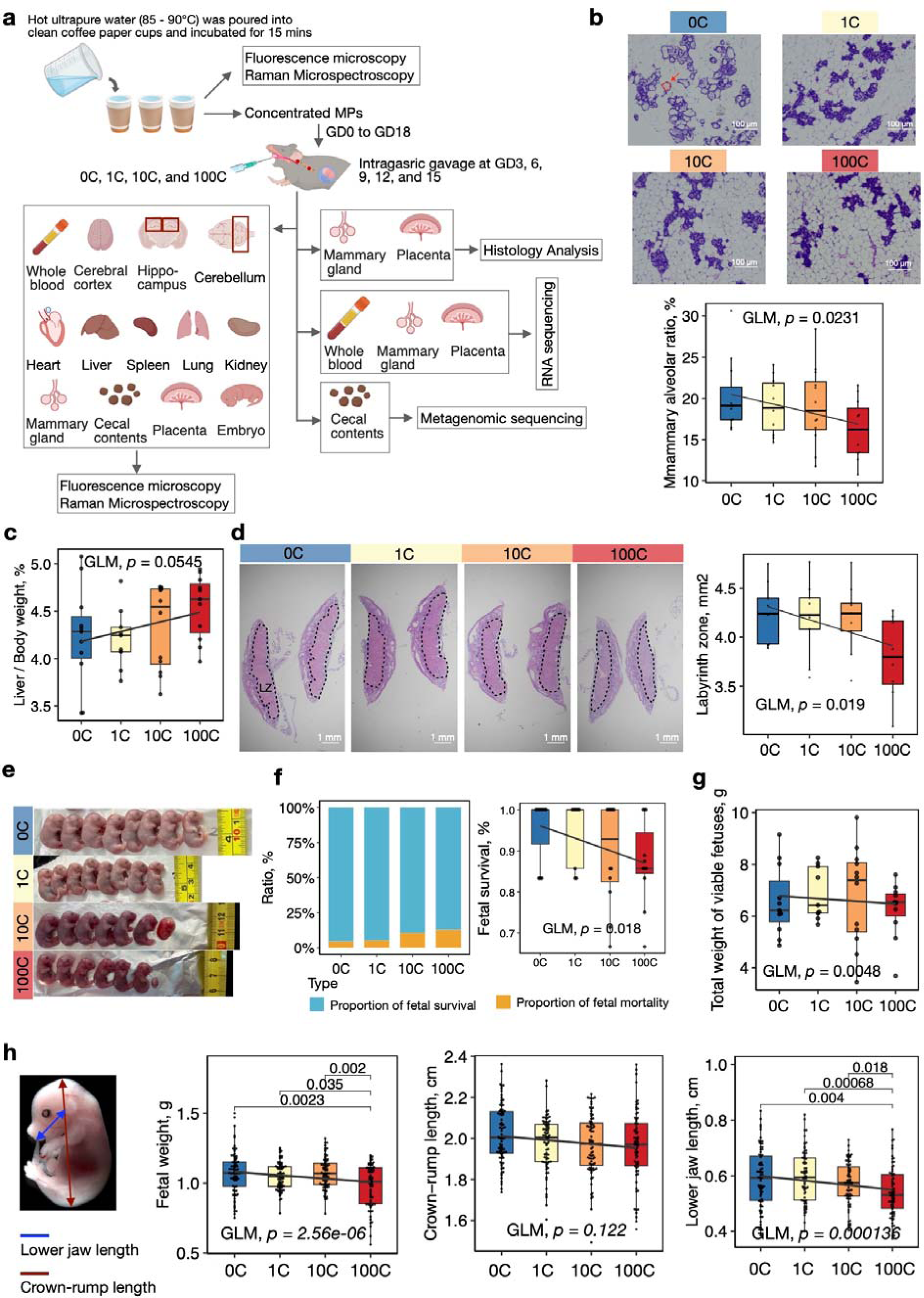
Top 5 enriched metabolic pathways. Metabolites with VIP >2 informed the metabolite set enrichment analysis of top 20 metabolites.

### JAS239 inhibits cell proliferation and blocks cell cycle progression, regardless of oxygenation, except in the 9L rat GBM cell line

We further aimed to probe the impact of JAS239 on cell proliferation under normoxic or hypoxic exposure. Based on the dose response curves (**Supplementary** Figure 2), we predicted JAS239 to exhibit similar effects on proliferation between normoxic and hypoxic conditions. We observed that F98 and 9L cells displayed a significant reduction in proliferation at 96 hours in the presence of JAS239 under normoxic conditions (**Figure 3 A & C**) without impacting cellular viability (**Figure 3 B and D**). Hypoxia only also impacted rat GBM cell proliferation in a comparable way to JAS239. There was no additive effect of the combination of the two conditions on F98, yet 9L cellular proliferation was further reduced in presence of both hypoxia and JAS239. In contrast, U-87 MG cell proliferation was not affected by hypoxic conditions (**Figure 3 E**). U-251 MG cells showed the strongest reduction in proliferation when treated with JAS239 in normoxia (**Figure 3 G**). JAS239 seemed slightly less effective in inhibiting proliferation of the hypoxic U-251 MG cells. In this case, the reduction in U-251 MG cell number over time is most likely due to induced cell death by 96 h of drug treatment given the significant reduction in cell viability (**Figure 3 F**). U-87 MG cell viability (**Figure 3 H**) was also reduced with JAS239 treatment by 96 h, however this was not significant.

**Figure 3.**
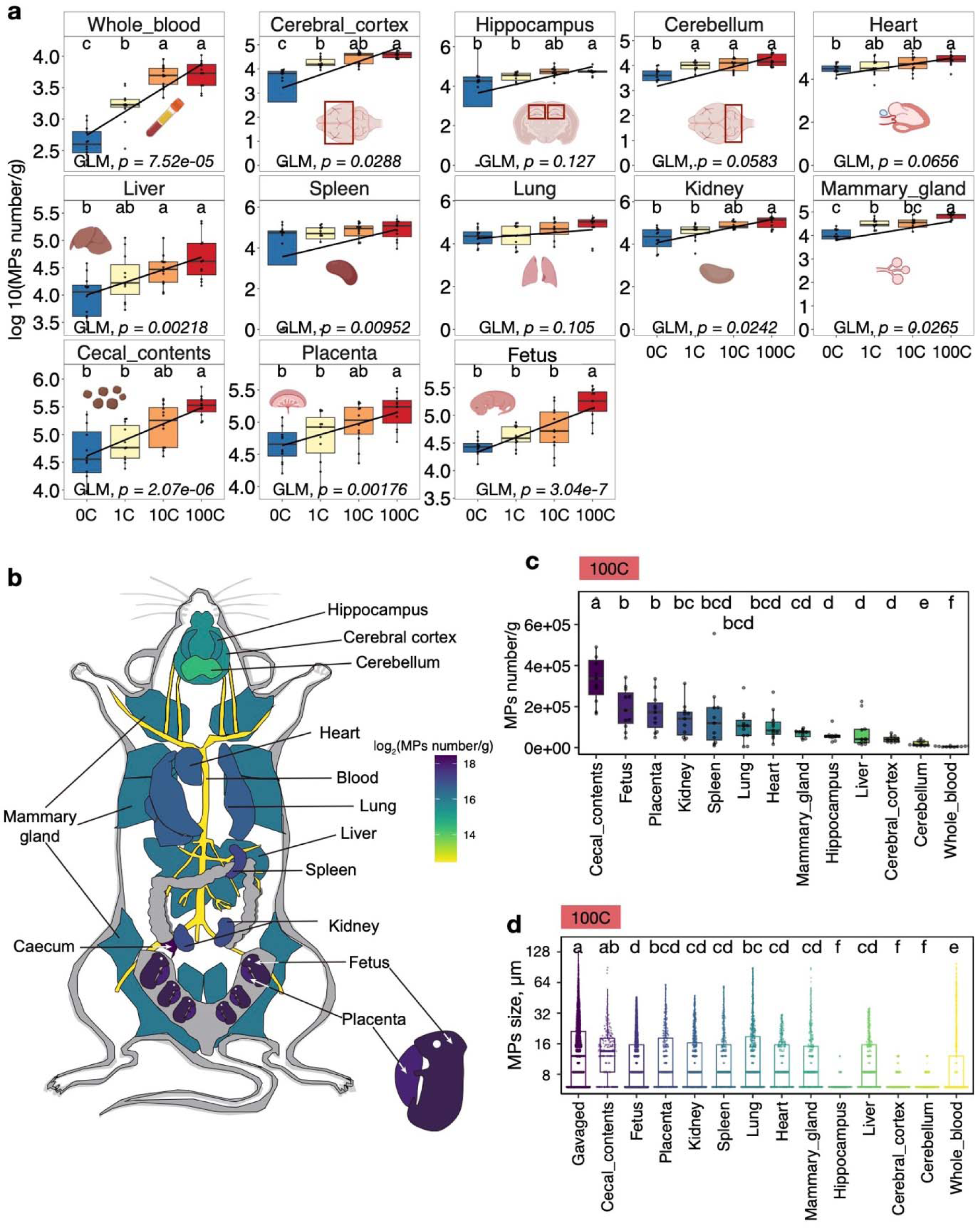
F98 (A, B), 9L (C, D), U-87 MG (E, F) and U-251 MG (G, H) cells were seeded and incubated in either 21% or 1% O_2_ and treated with either 500nM JAS239 or 0.1% DMSO control. Cell counts and viability were collected for each respective time point (n=3, +/- SEM). Multiple t-tests were performed on GraphPad Prism. Cell cycle distribution was acquired on a BD FACSCanto II system using a 405 laser and 440/50nm emission filter. Analysis was performed using Novo Express 1.4.1 software and cell cycle distributions were quantified. The histograms show mean values of three experiments for F98 (I), 9L (J), U87 (K) and U251 (L) cells. 2-way ANOVA of multiple comparisons was performed on GraphPad Prism. *p<0.05, **p<0.01, ***p<0.001, ****p<0.0001. Protein expression of E2F1 and E2F5 are shown for F98 (M), 9L (N), U-87 MG (O) and U-251 MG (P).

To further validate the proliferation data and to determine if the JAS239 induced reduction in proliferation was due to cell cycle arrest, we investigated the cell cycle distribution of GBM cells in response to JAS239 treatment. Cell cycle distribution was determined in viable, single cell populations using flow cytometry. We observed a significant increase in cells in G0/G1 in F98 (p<0.01, 72.3±11.5%), 9L (p<0.05, 63.4±11.3%) and U-251 MG (p<0.05, 22.6±5.9%) cells treated with JAS239 (**Figure 3 I, J & L**). Concomitantly, we also observed a significant reduction of cells in S phase for F98 (p<0.01, 19.6±8.7%) and U-87 MG cells (p<0.05, 10.8±2.6%; **Figure 3 I & K**) and a reduction of cells in G2/M phase for U-251 MG cells (p<0.05, 22.6±5.9%; **Figure 3 L**).

Similar to the proliferation data, the effects of JAS29 was comparable under normoxic and hypoxic conditions (**Figure 3 I, K & L**) in agreement with the MTT data (**Supplementary** Figure 2). The exception being 9L cells, where the combination of JAS239 and hypoxia had an enhancing effect on cell cycle arrest in G0/G1 with a significant increase in cells in G0/G1 between normoxic and hypoxic cells treated with JAS239 (63.4±11.2% versus 44.8, ±6.7% respectively, p<0.001). To depict the molecular mechanisms underlying the proliferation effects, we measured the protein expression levels of E2F transcription factors, E2F-1and 5. E2F-1 is a promoter of the G0/G1 to S phase transition and E2F-5 is considered as a repressor of cell cycle progression.[26] We observed a decrease of E2F-1 protein levels upon JAS239 treatment in all cell lines, regardless of oxygen levels (**Figure 3 M–P**), in line with the cell proliferation and cell cycle analysis above (**Figures 3**). Of note, the protein levels of the cell cycle repressor E2F-5 were comparable regardless of the conditions (JAS239 treatment and oxygen levels), with a notable exception being hypoxic JAS239-treated 9L cells, where a drastic increase in protein expression was observed. This observation is consistent with the cell cycle analysis results (**Figure 3 J**), where the most significant effect in these cells were observed with the hypoxia and JAS239 combination.

### Inhibition of cell invasion by JAS239 is cancelled by hypoxia

Whilst the effects of JAS239 on cell proliferation have been previously explored in different tumour types,[16, 18] we wanted to determine if JAS239 could also impact cell motility and invasive properties. We used a more physiological 3D spheroid model to assess invasion as it is difficult to assess this phenotype on cells grown as monolayers. The effects of JAS239 on cell invasion was determined in two GBM cell lines, the F98 (rat) and U87-MG (human). We first assessed the drug diffusion and distribution throughout the spheroid using the fluorescent properties of the drug. As the incubation system on the lightsheet microscope did not allow us to decrease oxygen levels, we used the hypoxia mimetic drug DMOG as a proxy for hypoxic conditions.

Similar to cells cultured in a 2D monolayer, JAS239 was localised in the cell cytoplasm (**Supplementary** Figure 8 **A–C**) yet its diffusion throughout all the cells within the 3D spheroid took approximately 77 min longer (80 min in the spheroid versus 3 min in cell monolayers, **Supplementary** Figure 8 **D & E**). JAS239 remained inside the cells for at least 48 h. A cell track analysis was then used to measure the position of the cell nuclei within the spheroid, and we determined typical mobility features including speed, track length, straightness, and displacement (dashed green line; **Figure 4 A**) over 16-h time-lapse experiments.

**Figure 4.**
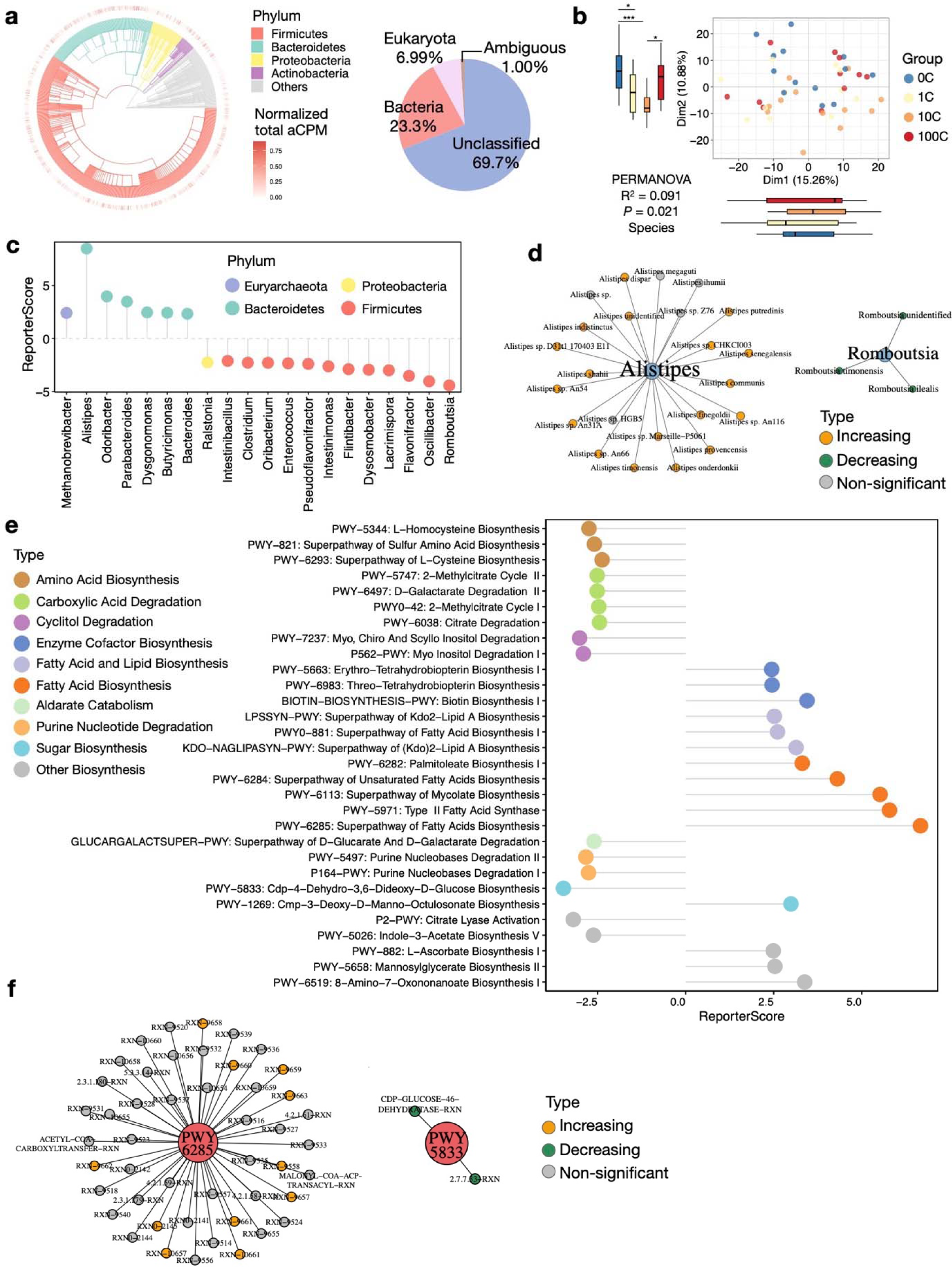
Schematic representation of the different types of cell movements and how each feature can be measured using Imaris software (A). The black lines show the potential directions and tracks that a cell could take from position A to position B. Track length is a measure of the black lines in these examples. The dashed green line represents the displacement of the cell, irrespective of the ‘distance’ the cell has moved. Track straightness and track speed can be determined using the track length. Spheroids were formed using 1.5x10^6^ F98 H2B RFP labelled cells and were incubated for 24 hours at 37°C/5% CO_2._ To mimic hypoxic conditioning, the prolyl hydroxylase inhibitor, DMOG (0.5mM) was added to the matrigel:media:hepes matrix during the spheroid mounting. For JAS239 treatment a final concentration of 500nM was applied to mounting media. Once mounted into FEP tubing, spheroids were left for 1 hour to immobilize before loading onto the Z.1 Light-Sheet microscope. A further 4 hours of setting time in the microscope chamber was allowed before the experiment started. Z-stacks of 6.5µM in thickness, every 3 minutes for 16 hours were acquired. (B) Representative spheroid pictures at time zero (T0) and 16 hours (T16) in both 21% an 1% O_2_, +/- 500nM JAS329. Using Imaris v6.9, tracks were filtered for minimum 2 hours in length. n=3. t-test with Welch’s correction was performed using GraphPad Prism.

The effect of JAS239 on 3D cell motility was visually striking on the time-lapse videos, as shown on the representative snapshots of F98 (**Figure 4 B**) and U-87 MG (**Figure 5 A**) spheroids. In comparison to untreated controls, less cells invaded the surrounding matrix after 16 h, and the spheroids appeared much more compact, indicating that JAS239 reduced invasive properties. To quantify this effect and assess cell displacement, we used mean squared displacement (MSD), a measure of average deviation of a particle from a reference point over time.[27] The type of diffusion can be determined from the pattern of the MSD trajectory as well as the value of the self-diffusion coefficient (D), as illustrated in **Figure 6 A**. The MSD plots for F98 and U-87 MG spheroids showed an increased slope with the addition of JAS239 in normoxic and DMOG conditions (**Figure 6 D & E**), indicating that JAS239 induced a more directed ‘super-diffusion’ motion in the cells, however, we postulate this displacement must be within the spheroid mass as opposed to the periphery of the spheroid, given that less cells invaded the surrounding matrix.

**Figure 5.**
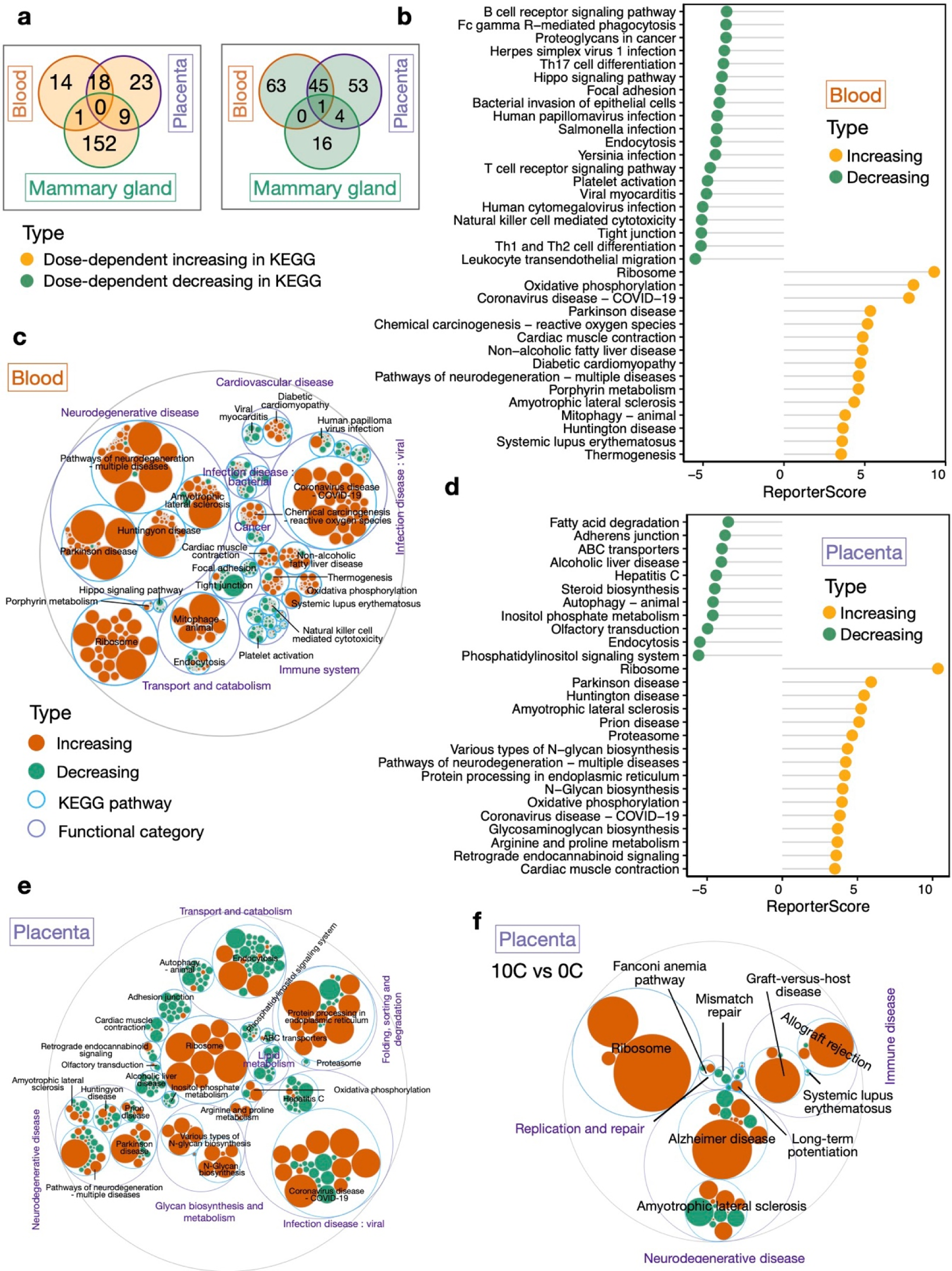
Spheroids were formed using 1.5x10^6^ U-87 MG H2B-RFP labelled cells and were incubated for 24 hours at 37°C/5% CO_2._ To achieve hypoxic conditioning, the prolyl hydroxylase inhibitor, DMOG (0.5mM) was added to Matrigel:media:hepes matrix upon spheroid mounting. For JAS239 treatment a final concentration of 500nM was applied to mounting media. Once mounted into FEP tubing, spheroids could set for 1 hour before loading onto the Light-Sheet microscope and a further 4 hours on the microscope before the experiment started. Z-stacks of 6.5µM in thickness, every 3 minutes for 16 hours were acquired. Screen shots (A) of representative spheroid pictures at time zero (T0) and 16 hours (T16) in both 21% an 1% O_2_, +/- 500nM JAS329. Using Imaris v6.9, tracks were filtered for minimum 2 hours in length. n=3. t-test with Welch’s correction was performed using Graphpad Prism.

**Figure 6.**
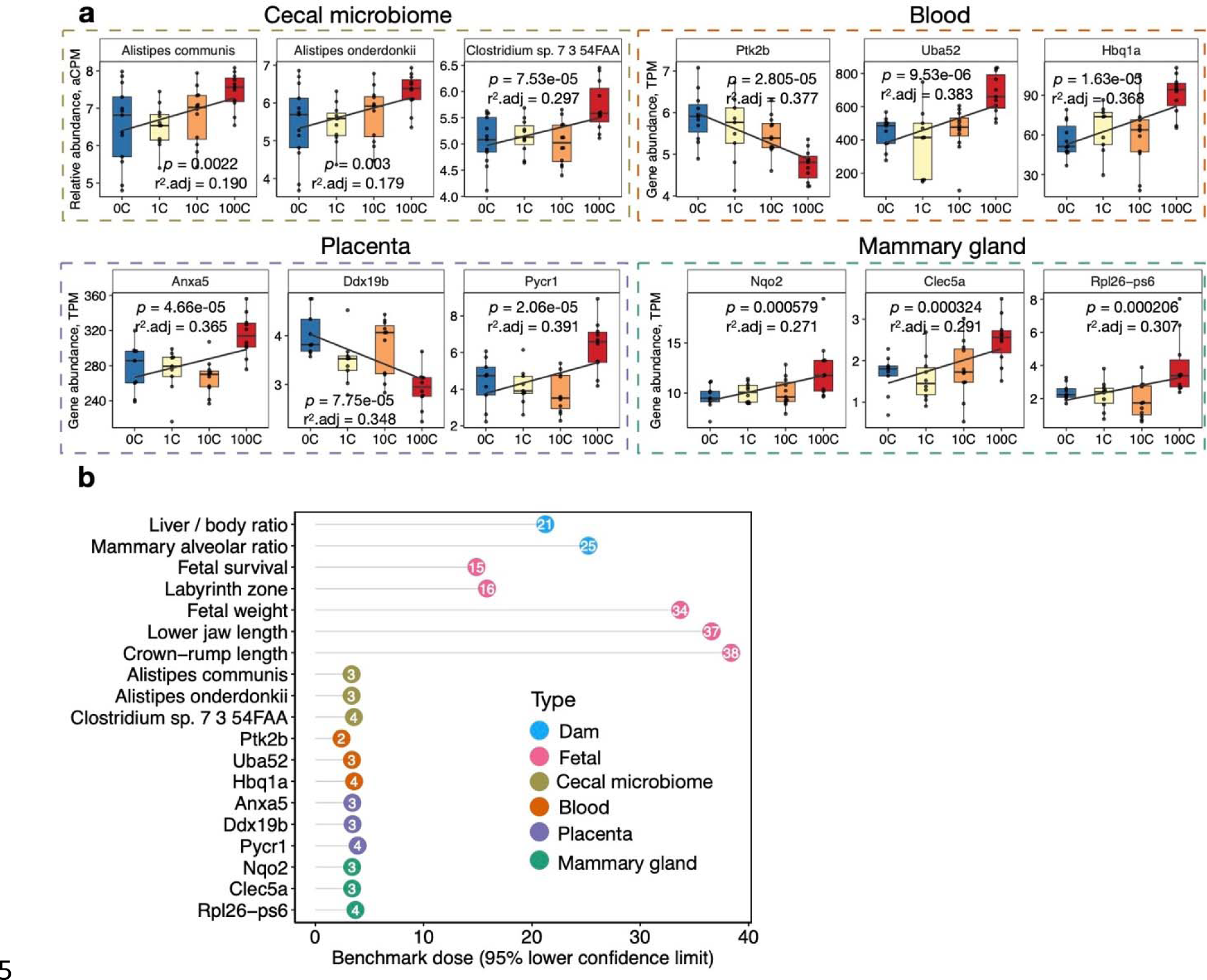
Schematic representation of the MSD (R^2^) versus time for diffusion and flow, normal diffusion, sub-diffusion, and confined diffusion (A). D, self-diffusion coefficient; figure adapted from Saxton and Jacobson, 1997. Mean Squared Displacement (MSD; R^2^) analysis of F98 (B & D) & U-87 MG (C & E) H2B RFP labelled spheroids were assessed using Imaris v6.9 for tracks at least 2 hours in length. Dot plots of combined MSD are shown for F98 (A) and U-87 MG spheroids in control and DMOG treated (0.5 mM) with 500 nM JAS239. MSD trajectories for F98 (D) and U-87 MG (E) spheroids are also shown +/- SEM, n=3. MSD, mean squared displacement.

To further evaluate overall cell movement in the F98 and U-87 MG spheroids, we measured several features including track speed, straightness and length, which, combined, are more informative of the cells’ behavior. JAS239 significantly reduced track length in F98 spheroids compared to untreated controls (113.2±86 µm versus 84.0±48.0 µm; p<0.0001, **Figure 4 D**), but in contrast, induced an increase in track length in U-87 MG spheroids (120.7±57.7 µm versus 133.0±61.6 µm; p<0.001 **Figure 5 C**). Track straightness provides better information on the invasiveness through division of displacement over track length. A track straightness value closer to 0 infers more disorganized movement whereas a value closer to 1 implies a straighter path. F98 cells showed a significant reduction in track straightness with JAS239 treatment (control: 0.32±0.12; JAS239: 0.28±0.11; p<0.0001, **Figure 4 C**) whereas no changes in straightness was observed for U-87 MG (**Figure 5 B**). JAS239 significantly reduced the average speed of cells in F98 spheroids (control: 0.007±0.004 µm/sec, JAS239: 0.005±0.001 µm/sec; p<0.0001, **Figure 4 E**), however the opposite was observed in U-87 MG JAS239 treated spheroids (control: 0.006±0.001 µm/sec; JAS239: 0.007± 0.001 µm/sec, p<0.0001, **Figure 5 D**). It therefore suggests that JAS239 affects cell motility in a cell-line-dependent manner. Next, we wanted to determine the impact of hypoxia on JAS239 treatment on these cell motility parameters. Compared to controls, addition of DMOG significantly reduced F98 and U87 track straightness and increased track length and speed (**Figure 4 C–E** p<0.0001 and **Figure 5 B–D** p<0.05, respectively). Addition of JAS239 to DMOG treatment significantly reduced F98 cell track straightness, length and, surprisingly, increased cell speed (**Figure 4 C–E**; p<0.0001). Track straightness and speed in DMOG and JAS239 treated U-87 MG cells were comparable to DMOG only (**Figure 5 B & D**), however, similar to F98 spheroids, there was a significant increase in track speed in presence of both DMOG and JAS239 (**Figure 5 D**).

Collectively, these data suggest JAS239 impacts key invasive properties and reduces the invasive potential of GBM cells into the surrounding matrix, and its effects vary between control and DMOG conditions. In F98 spheroids, DMOG & JAS239 treatment induced faster, more super-diffusive motion than JAS239 only spheroids but both conditions induced a disorganized cell movement, with significantly shorter track lengths. Similarly, in U-87 MG spheroids, DMOG & JAS239 treatment induced faster, more super-diffusive motion than spheroids treated with only JAS239.

To better explain the invasion parameters, we assessed the protein expression of integrin α3β1 as a marker of cell mobility as α3β1 mediates migration of neuronal cells.[28] We observed reduced α3β1 levels in F98 (**Supplementary** Figure 9 **A**), 9L (**Supplementary** Figure 9 **B**) and U- 251 MG (**Supplementary** Figure 9 **D**) cells treated with JAS239, in line with the decreased invasion observed in the spheroids. Yet, U-87 MG showed no change in α3β1 expression (**Supplementary** Figure 9 **C**), suggesting that other cell migration signalling molecules may be involved in the reduction in cell invasion previously observed upon JAS239 treatment (**Figure 5**).

## Discussion

We explored the effects of the hypoxic microenvironment found in tumours, on the GBM cell response to choline kinase inhibition. Beyond the classical effects expected from drugs affecting cellular metabolism (cell proliferation, survival, cell cycle progression), we also assessed cell invasion. Interestingly, the choline kinase inhibitor JAS239, typically assessed for its antiproliferative effects in tumour cells, also demonstrated clear inhibitory effects on cell invasion. This observation along with the fact that JAS239 potency was not damped by hypoxic conditions, to which tumour cells are exposed, makes this drug an interesting candidate for GBM treatment as it has the potential to not only reduce tumour growth but also its invasion, which remains one of the major clinical challenges in the treatment of GBM. Additionally, our work on different GBM cell lines shows an important cell line to cell line variability, which must be considered for future potential use in the clinic.

### JAS239

The effects of JAS239 have never been explored in hypoxic tumour cells. Under normoxic conditions, JAS239 induced a significant reduction in proliferation via a cell cycle arrest in all cell lines, which was expected, given its effects in reducing phosphatidylcholine synthesis used for cell membrane synthesis. Under hypoxic conditions, we observed no synergistic effect of JAS239 on proliferation or cell cycle arrest. Yet, we found JAS239 to be equally efficacious under hypoxic conditions with respect to reducing cell invasion, which given the high mortality of GBM being largely attributed to its highly invasive nature, makes it a promising treatment strategy for GBM patients.

The effects of other choline kinase inhibitors on cancer cell migration and invasion were previously observed. EB-3D-mediated choline kinase inhibition showed a significant reduction in the migratory and invasive properties of the highly aggressive MDA-MB-231 breast cancer cell line.[29] Although these cell invasion assays were carried out in 2D trans-well plate inserts, these data are in line with what we observed in highly aggressive GBM lines using JAS239 in 3D spheroid models. Interestingly Mariotto *et al*, also demonstrated reduced *in vivo* lung metastases with inhibition of choline kinase, providing further evidence of the potential for choline kinase inhibitors as effective treatments for aggressive, highly invasive cancers.[29]

However, these drugs will probably reach their full promise when used in combination with other therapies. For example, inhibition of ChoKα has been shown to increase the sensitivity of aggressive endothelial ovarian cancer cells to commonly used chemotherapeutics including paclitaxel and doxorubicin in breast cancer cells.[30, 31]

The efficacy of JAS239 treatment has also been demonstrated in *in vivo* rodent models of breast cancer and GBM.[16, 32] The initial studies by Arlauckas *et al*, reported JAS239 efficacy with a significant reduction in total choline levels and tumour growth in a breast cancer xenograft model.[16] Additionally, in previous *in vivo* experiments we used ^1^H NMR spectroscopy (^1^H MRS) to assess JAS239 efficacy in three GBM xenograft rodent models (GL261, F98 and 9L) and demonstrated a reduction in total choline levels in all three models in response to JAS239, with the 9L intracranial tumours exhibiting the largest reduction. [32] Interestingly, the opposite trend was observed in tumour volume; when compared to baseline, the largest tumour growth arrest was found in GL261 and F98 tumours followed by 9L tumours.[32] These *in vivo* data support observations we have made in this study with respects to JAS239 induced reductions in proliferation via cell cycle arrest, with no effect on cell viability. The *in vitro* data obtained in this study, also explain why *in vivo* conditions, only a tumour growth arrest was observed, as opposed to a reduction in tumour volume [21]. Moreover, the largest reduction in choline levels occurred within the tumour region and this coincided with reductions in mitotic index for all three GBM models, suggestive that JAS239 selectively inhibits proliferating cells within the tumour, where choline kinase is over expressed.

### Hypoxia and GBM

Hypoxia is an important hallmark of GBM and is attributed to its highly aggressive and invasive phenotype. Adaptation to hypoxia is mediated via HIF-1/2α and typically involves the reduction of high energy consuming pathways, such as cell division.[13, 33] How cells respond to hypoxia can be dependent on the severity and duration of hypoxia conditioning, however most cells respond via arresting at G1 or early S phase of the cell cycle.[34] The exact mechanism is disputed across the literature as some studies suggest the G1/S arrest is due to hypoxia induced increase in p27 levels.[35] Contrary to this, a study investigating the effects of hypoxia on cell cycle response across multiple cell lines reported only one cell line exhibiting hypoxia-induced increase in p27 mRNA and two at the protein level. HeLa cells had a reduction in p27 in response to hypoxia yet arrested in G1/S phase.[34] The authors did however note the cell type dependent response to hypoxia. Our data are very similar to previous observations by Richards *et al*, where a panel of human GBM cell lines (D566, U-87 MG and U-251 MG) were assessed in varying oxygen concentrations (0.1% O_2_ and 1% O_2_) for 24–72 hours, with no change in cell cycle distribution even at 0.1% O_2_. No change in expression levels of p21, p27 and G1/S transition transcription factor E2F1 were reported in that study.[33] Our data confirm the cell type dependent response to hypoxia. Indeed, out of the four cell lines tested, only F98 cells demonstrated a significant G0/G1 cell cycle arrest after 96 hours of hypoxia conditioning; the other cell lines appeared to have more cells in G2/M phase, yet this was not significant. Overall, the rat GBM cells appeared more sensitive to prolonged hypoxia compared to both human lines, however the U-87 MG cells were notably the least sensitive of all cells. Since a cell adaptation to hypoxia is mediated via HIF- 1/2α, we assessed the protein stability of HIF-1/2α over time. Of the cell lines studied, the U-87 MG cells had the most HIF-1α stabilization between 24–72 hours while U-251 MG cells showed consistent HIF-1α stabilization even under normoxic conditions. The differences in HIF-1α protein expression over time may explain the differences the cell specific sensitivity to hypoxia observed.

Another potential mechanism by which hypoxia induces pro-tumorigenic effects is through its enhancement of tumour invasion.[36, 37] In a study assessing U-87 MG and U-251 MG migration and invasion, cells conditioned for 72 hours in hypoxia (1% O_2_) demonstrated drastic differences in migration potential. The increase in migration potential of U-251 MG cells was limited compared to U-87 MG cells, where migration doubled in a wound healing assay.[38] In our U-87 MG spheroid invasion experiments, we observed a significant increase in track straightness, length and speed between normoxia and DMOG-hypoxia. Additionally, we also observed an increase in MSD under DMOG treatment, indicative of super-diffusion motion. Joseph *et al* correlated this hypoxia-induced increase in migratory potential of cells with a mesenchymal transition mediated by HIF-1α and ZEB1.[38] GBMs with a mesenchymal signature are suggested to be of particular importance as they are deemed to be highly aggressive compared to the other subtypes, proneural, neural and classical and are considered to be highly resistant to therapies;[39, 40] U-87 MG cells are considered to be a model for the mesenchymal subtype of GBM.

### JAS239 and hypoxia

An unexpected but promising finding of this study was the JAS239 induced reduction of HIF- 1/2α protein levels across all cell lines. Given the vast number of pro-tumourigenic genes regulated by HIF transcription factors and the association with treatment resistance [41, 42], this indirect effect of JAS239 adds to the potential of this inhibitor in hypoxic tumours. Several direct and indirect HIF inhibitors (panzem, tanespimycin, zolinza) have been tested in phase I/II clinical trials of multiple myeloma, ovarian cancer, prostate cancer and lymphoma, however these trials have not reported significant improvement in overall survival and modest progression free survival compared to current standard of care.[43] The potential of JAS239 as an inhibitor of HIF, in addition to its metabolic, anti-proliferative and anti-invasive properties, warrants further investigation and could serve as a candidate treatment in combination with standard of care treatments in GBM to target multiple aspects of tumourigenesis.

## Conclusion

Whilst we previously demonstrated anti-tumour activity of JA239 *in vivo*,[32] we here provide further understanding of its molecular mechanisms of action including the changes in intracellular metabolites and the cellular effects on proliferation and invasion. We noticed an important cell line to cell line variability, likely representative of the clinical heterogeneity of these tumours displaying a range of molecular subtypes and mutational burden. As a consequence of this heterogeneity, human GBMs have been categorized into defined molecular subgroups. Our findings about the ability of a single drug (JAS239) to obliterate at once, at least 3 different hallmarks of cancer and one enabling characteristic:[44] unlimited proliferation, deregulating cellular metabolism, activating invasion, activation of HIF; opens the way for new promising strategies that can act on multiple pro-tumorigenic features, increasing the chance to better eradicate tumour development in patients.

## List of Abbreviations

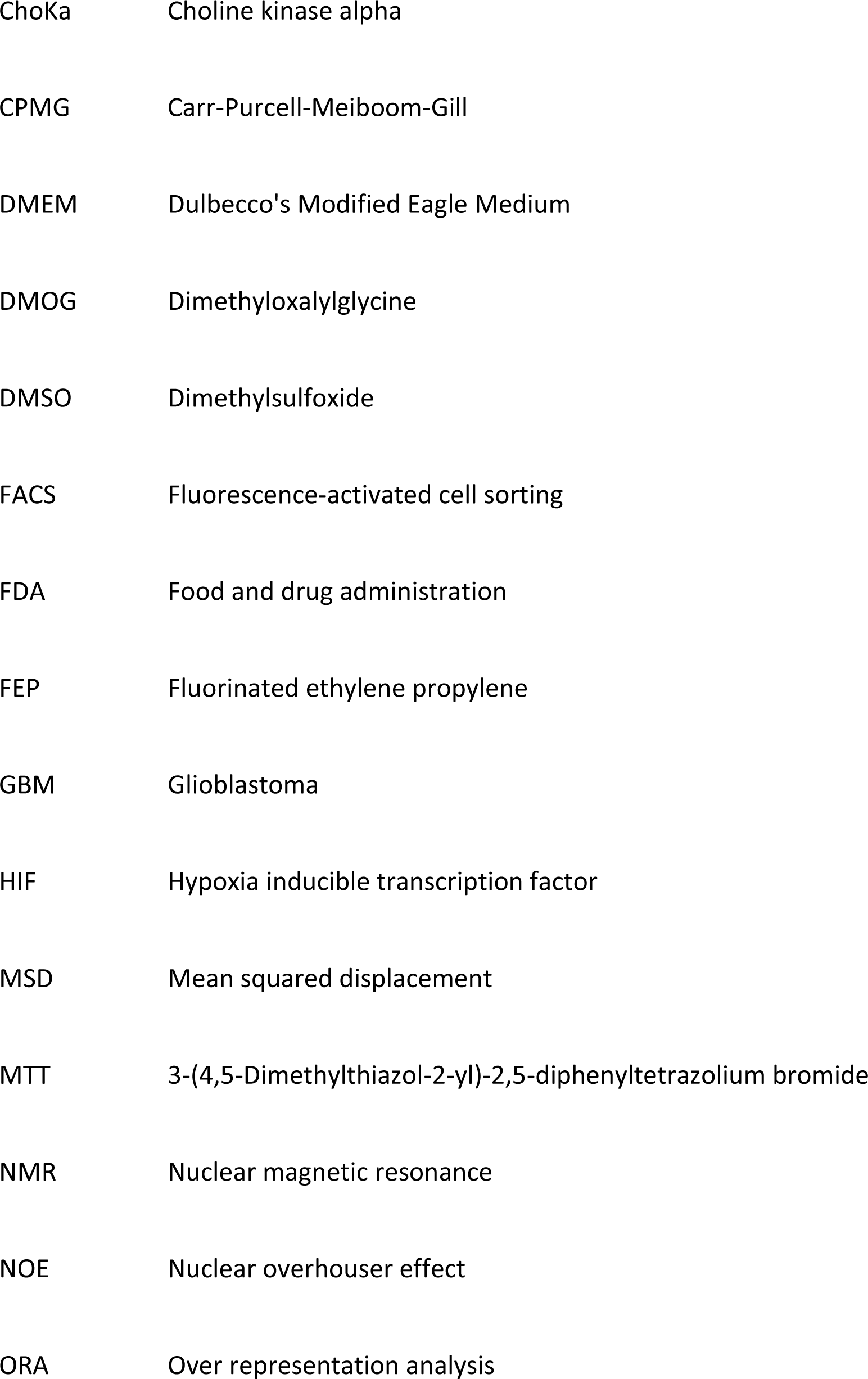
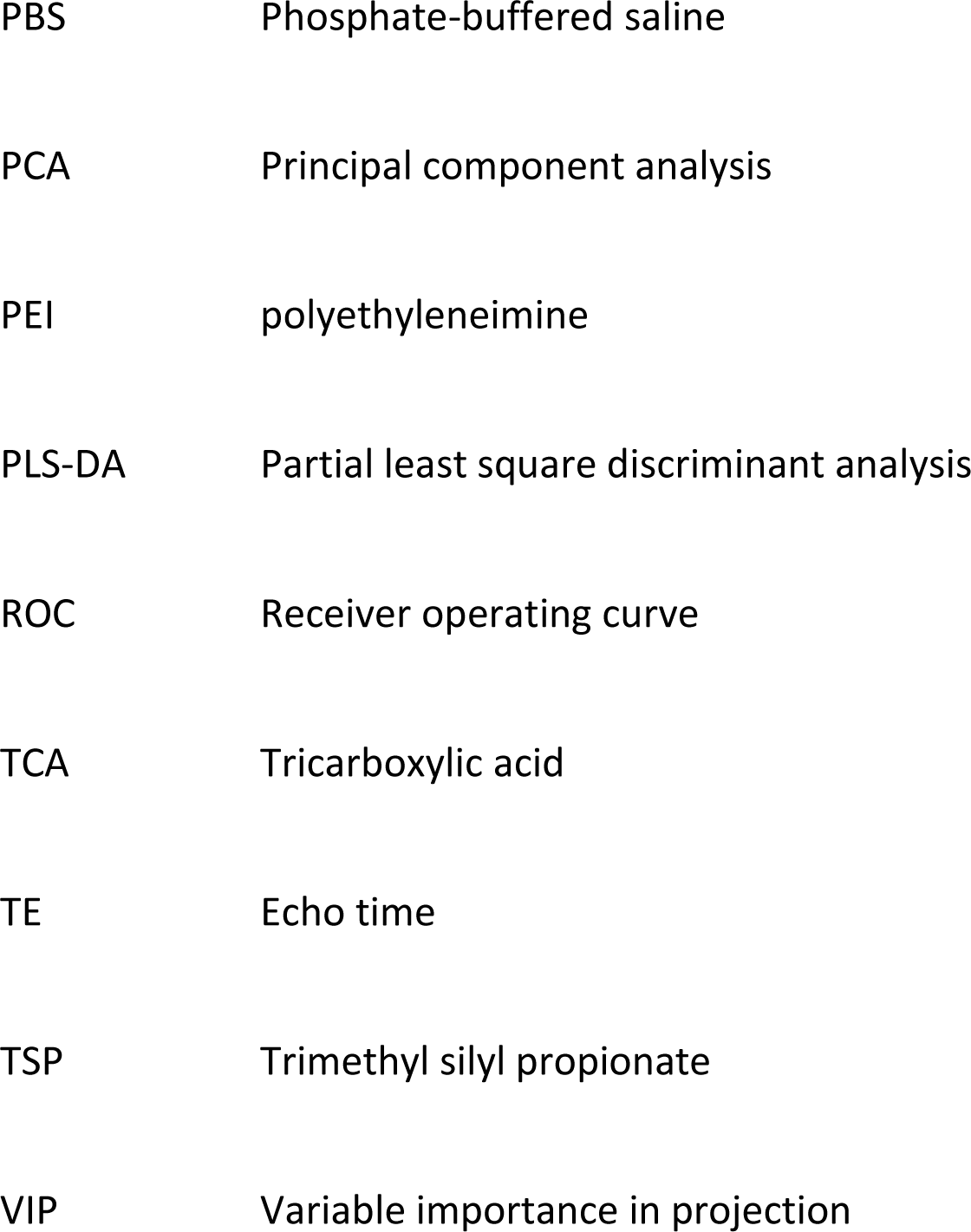

## Declarations

### Ethics approval and consent to participate

Not applicable

### Consent for publication

Not applicable

### Availability of data and materials

The raw data files presented in this study are available on request from the corresponding authors. Additionally, the NMR data files from the current study are available in www.ebi.ac.uk/MetaboLights ID MTBLS 6212

### Competing Interests

The authors have no competing interests to declare.

### Funding

No external funding sources were involved in the design of the study; collection, analyses, data interpretation, writing of the manuscript, or decision to publish the results.

### Author’s contributions

CLK maintained the cell lines, performed all hypoxia and treatment studies, performed cell extracts. CLK and MW performed the 3D spheroid experiments. CLK and MP performed the NMR metabolomic studies as well as generation of respective figures. SF and EJD synthesised and provided JAS239 and also helped with in vitro assays. CLK, VS, HP were involved in the design of the experiments, data analysis, and generation of figures. CLK, EJD, VS and HP contributed to manuscript writing, generation of figures and interpretation of data. All authors read and approved the final manuscript.

## Acknowledgments

The authors acknowledge the use of the Centre for Cell Imaging (CCI; Zeiss Lightsheet Z.1 was funded by BBSRC Alert13 grant number <u>BB/L014947/1</u>) and the Highfield NMR Facilities provided by Liverpool Shared Research Facilities, Faculty of Health and Life sciences, University of Liverpool. We would especially like to thank Jennifer Adcott and Marie Held from the CCI for their precious time and help in microscopy and image data analysis. We would also like to thank Dr Thomas Leather for assistance with NMR analysis and Raven Chandramohan and Daniel Thomas of Ngee An Polytechnic for assistance with NMR sample preparation. Claire L Kelly was funded by a PhD studentship from the Northwest Cancer Charity. Dr Anne Hermann of the University of Liverpool is acknowledged for her assistance with production of F98 and U-87 MG H2B RFP cells. Dr Arthur Taylor of the University of Liverpool is acknowledged for his assistance with production of F98-Luc-ZS Green and 9L-Luc-ZS Green cells. Dr Delikatny acknowledges the support of the National Cancer Institute R01 CA226412.

## Notes

### Competing Interest Statement

The authors have declared no competing interest.

## References

1. Li, S., et al., Signaling pathways in brain tumors and therapeutic interventions. Signal Transduct Target Ther, 2023. 8(1): p. 8.

2. Nagy, A., et al., Molecular Subgroups of Glioblastoma- an Assessment by Immunohistochemical Markers. Pathol Oncol Res, 2019. 25(1): p. 21–31.

3. Davis, M.E., Glioblastoma: Overview of Disease and Treatment. Clin J Oncol Nurs, 2016. 20(5 Suppl): p. S2-8.

4. Society;, A.C. Survival Rates for Selected Adult Brain and Spinal Cord Tumors. 2022 [cited 2023 12th March]; Available from: https://www.cancer.org/cancer/brain-spinal-cord-tumors-adults/detection-diagnosis-staging/survival-rates.html.

5. UK;, C.R. *Brain Tumours: Survival*. 2022 [cited 2023 12th March]; Available from: https://www.cancerresearchuk.org/about-cancer/brain-tumours/survival.

6. Cohen, M.H., et al., FDA drug approval summary: bevacizumab (Avastin) as treatment of recurrent glioblastoma multiforme. Oncologist, 2009. 14(11): p. 1131–8.

7. Friedman, H.S., T. Kerby, and H. Calvert, Temozolomide and treatment of malignant glioma. Clin Cancer Res, 2000. 6(7): p. 2585–97.

8. Huang, B., et al., Advances in Immunotherapy for Glioblastoma Multiforme. J Immunol Res, 2017. 2017: p. 3597613.

9. Boyd, N.H., et al., Glioma stem cells and their roles within the hypoxic tumor microenvironment. Theranostics, 2021. 11(2): p. 665–683.

10. Hambardzumyan, D. and G. Bergers, Glioblastoma: Defining Tumor Niches. Trends Cancer, 2015. 1(4): p. 252–265.

11. Hambardzumyan, D., D.H. Gutmann, and H. Kettenmann, The role of microglia and macrophages in glioma maintenance and progression. Nat Neurosci, 2016. 19(1): p. 20–7.

12. Jamal, M., et al., The brain microenvironment preferentially enhances the radioresistance of CD133(+) glioblastoma stem-like cells. Neoplasia, 2012. 14(2): p. 150–8.

13. Ortmann, B., J. Druker, and S. Rocha, Cell cycle progression in response to oxygen levels. Cell Mol Life Sci, 2014. 71(18): p. 3569–82.

14. Wang, P., et al., HIF1alpha regulates single differentiated glioma cell dedifferentiation to stem-like cell phenotypes with high tumorigenic potential under hypoxia. Oncotarget, 2017. 8(17): p. 28074–28092.

15. Sobanski, T., et al., Cell Metabolism and DNA Repair Pathways: Implications for Cancer Therapy. Front Cell Dev Biol, 2021. 9: p. 633305.

16. Arlauckas, S.P., A.V. Popov, and E.J. Delikatny, Direct inhibition of choline kinase by a near-infrared fluorescent carbocyanine. Mol Cancer Ther, 2014. 13(9): p. 2149–58.

17. Arlauckas, S.P., A.V. Popov, and E.J. Delikatny, Choline kinase alpha-Putting the ChoK-hold on tumor metabolism. Prog Lipid Res, 2016. 63: p. 28–40.

18. Arlauckas, S.P., et al., Near infrared fluorescent imaging of choline kinase alpha expression and inhibition in breast tumors. Oncotarget, 2017. 8(10): p. 16518–16530.

19. Kumar, M.A., S.P; Saksena, S; Verma, G; Ittyerah, R; Pickup, S; Popov, A.V; Delikatny, E.J; Poptani, H;, Magnetic Resonance Spectroscopy for Detection of Choline Kinase Inhibition in the Treatment of Brain Tumors. Molecular Cancer Therapeutics 2015. 14(1): p. 899–908.

20. Trousil, S., et al., The novel choline kinase inhibitor ICL-CCIC-0019 reprograms cellular metabolism and inhibits cancer cell growth. Oncotarget, 2016. 7(24): p. 37103–37120.

21. Pang, Z., et al., MetaboAnalyst 5.0: narrowing the gap between raw spectra and functional insights. Nucleic Acids Res, 2021. 49(W1): p. W388–W396.

22. Li, Y., L. Zhao, and X.F. Li, The Hypoxia-Activated Prodrug TH-302: Exploiting Hypoxia in Cancer Therapy. Front Pharmacol, 2021. 12: p. 636892.

23. Strese, S., et al., Effects of hypoxia on human cancer cell line chemosensitivity. BMC Cancer, 2013. 13: p. 331.

24. Zhu, J., et al., High-altitude Hypoxia Influences the Activities of the Drug-Metabolizing Enzyme CYP3A1 and the Pharmacokinetics of Four Cardiovascular System Drugs. Pharmaceuticals (Basel), 2022. 15(10).

25. Cowman, S., Elucidation of the Effects of Hypoxia on DNA Repair Machinery in Brain Tumour Cells, in Institute of Systems, Molecular and Integrative Biology. 2018, University of Liverpool: Livrepository. p. 288.

26. Bertoli, C., J.M. Skotheim, and R.A. de Bruin, Control of cell cycle transcription during G1 and S phases. Nat Rev Mol Cell Biol, 2013. 14(8): p. 518–28.

27. Saxton, M.J. and K. Jacobson, Single-particle tracking: applications to membrane dynamics. Annu Rev Biophys Biomol Struct, 1997. 26: p. 373–99.

28. Nakada, M., et al., Integrin alpha3 is overexpressed in glioma stem-like cells and promotes invasion. Br J Cancer, 2013. 108(12): p. 2516–24.

29. Mariotto, E., et al., Choline Kinase Alpha Inhibition by EB-3D Triggers Cellular Senescence, Reduces Tumor Growth and Metastatic Dissemination in Breast Cancer. Cancers (Basel), 2018. 10(10).

30. Granata, A., et al., Choline kinase-alpha by regulating cell aggressiveness and drug sensitivity is a potential druggable target for ovarian cancer. Br J Cancer, 2014. 110(2): p. 330–40.

31. Mori, N., et al., Choline kinase down-regulation increases the effect of 5-fluorouracil in breast cancer cells. Cancer Res, 2007. 67(23): p. 11284–90.

32. Bhaduri, S., et al., Assessing Tumour Haemodynamic Heterogeneity and Response to Choline Kinase Inhibition Using Clustered Dynamic Contrast Enhanced MRI Parameters in Rodent Models of Glioblastoma. Cancers (Basel), 2022. 14(5).

33. Richards, R., et al., Cell cycle progression in glioblastoma cells is unaffected by pathophysiological levels of hypoxia. PeerJ, 2016. 4: p. e1755.

34. Box, A.H. and D.J. Demetrick, Cell cycle kinase inhibitor expression and hypoxia-induced cell cycle arrest in human cancer cell lines. Carcinogenesis, 2004. 25(12): p. 2325–35.

35. Gardner, L.B., et al., Hypoxia inhibits G1/S transition through regulation of p27 expression. J Biol Chem, 2001. 276(11): p. 7919–26.

36. Fujiwara, S., et al., Silencing hypoxia-inducible factor-1alpha inhibits cell migration and invasion under hypoxic environment in malignant gliomas. Int J Oncol, 2007. 30(4): p. 793–802.

37. Hoffmann, C., et al., Hypoxia promotes breast cancer cell invasion through HIF-1alpha-mediated up-regulation of the invadopodial actin bundling protein CSRP2. Sci Rep, 2018. 8(1): p. 10191.

38. Joseph, J.V., et al., Hypoxia enhances migration and invasion in glioblastoma by promoting a mesenchymal shift mediated by the HIF1alpha-ZEB1 axis. Cancer Lett, 2015. 359(1): p. 107–16.

39. Bhat, K.P.L., et al., Mesenchymal differentiation mediated by NF-kappaB promotes radiation resistance in glioblastoma. Cancer Cell, 2013. 24(3): p. 331–46.

40. Mikheeva, S.A., et al., TWIST1 promotes invasion through mesenchymal change in human glioblastoma. Mol Cancer, 2010. 9: p. 194.

41. Strowd, R.E.E., B.M; Wen, P.Y; Ahluwalia, M.S; Piotrowski, A.F; Desai, A,S; . *Safety and activity of a first-in-class oral HIF2-alpha inhibitor, PT2385, in patients with first recurrent glioblastoma (GBM).* in *ASCO*. 2019. Journal of Clinical Oncology.

42. Wang, E., et al., The role of factor inhibiting HIF (FIH-1) in inhibiting HIF-1 transcriptional activity in glioblastoma multiforme. PLoS One, 2014. 9(1): p. e86102.

43. Fallah, J. and B.I. Rini, HIF Inhibitors: Status of Current Clinical Development. Curr Oncol Rep, 2019. 21(1): p. 6.

44. Hanahan, D., Hallmarks of Cancer: New Dimensions. Cancer Discov, 2022. 12(1): p. 31–46.

