## supplementary text and figures for "Inhibition of glioblastoma cell proliferation and invasion by the choline-kinase inhibitor JAS239 varies with cell type and hypoxia"

### Supplementary Section

#### Supplementary Methods Text

##### Section 1

***Cell line information***

The rat F98 (source American Type Culture Collection [ATCC] CRL-2397) and 9L LacZ (source ATCC CRL-2200) GBM cells were cultured in Gibco Dulbecco's Modified Eagle Medium high glucose (4.5 g/L) medium supplemented with 1% L-Glutamine and 10% foetal bovine serum (FBS). Human Uppsala 87 malignant glioma (U-87 MG; source ATCC HTB-14 TM) and Uppsala 251 malignant glioma (U-251 MG; source Cell Line Services Eppelheim, Germany) cells were cultured in Gibco Minimum Essential Medium supplemented with 1% sodium pyruvate and 10% FBS.

***Stable cell line production***

A lentiviral transduction method was used to insert the pHIV-H2BmRFP plasmid (addgene plasmid number #18982) or pHIV-Luc-ZsGreen (addgene plasmid number #39196) plasmid. Briefly, 1.5x106 HEK293T cells were seeded per 10cm dish and incubated overnight at 37°C. Cells were transfected with the lentiviral reagent, made up in a 4:2:1 ratio of vector:packaging:envelope. The plasmids were then transfected with polyethylenimine (PEI) at a ratio of 2:1 PEI:DNA in serum free DMEM medium and incubated at 37°C for maximum of 16 h, followed by a media change. Three days after transfection, the conditioned media was collected, centrifuged at 1000 g for 5 minutes (min) and filtered through a 0.45 µM polyethersulfone filter. Ultracentrifugation was then carried out according to the protocol described by Kutner et al.[1] Virus containing media was transferred to ultracentrifugation tubes, a 20% sucrose cushion was expelled beneath the media layer and then ultracentrifugation was performed at 4°C for 2 h at 21,000 rpm. Viral particles were concentrated in 600 µl PBS and applied to F98 and U-87 MG cells immediately. For transduction, 1.4x105 F98 and U-87 MG cells were seeded into a T25 flask. The full 600 µl of concentrated virus was applied and a media change was performed after 72 h.

***Cell viability***

Cells were preincubated for 72 h at 21% $O_2$ or 1% $O_2$ at 37°C and then treated with JAS239 using a half-log serial dilution from 100 µM. After treatment, cells were further incubated for 24 h in the same incubation conditions. Following treatment, 5 µl of 5 mg/ml MTT reagent (Sigma Aldrich) was added and incubated for 4 h. 50 µl of 10% SDS /0.01 M HCl solubilizing reagent was added and incubated overnight at 21% $O_2$/37°C. Absorbance was read at 570 nm using a Spectramax plus384 (Molecular Devices) plate reader.

***NMR***

*Acetonitrile Extraction*

Aqueous metabolites were extracted using 400 µl of ice-cold solution of 50% HPLC-grade acetonitrile: 50% double distilled water per sample. Samples were sonicated in an ice bath (4°C) for 3x 30 seconds (sec) at 10 kHz using a microtip probe. Samples were then vortexed and centrifuged for 5 min/12,000 xg/ 4°C. The cell pellets were discarded and the supernatants were snap frozen in liquid nitrogen and lyophilized overnight. Samples were stored at -80°C until further use.

*1H NMR Sample Preparation and data Acquisition*

Samples were reconstituted in of a solution containing sodium phosphate (pH 7.4), deuterated (d4) trimethyl silyl propionate (TSP), 1.2 mM sodium azide and 99.8% 2H2O. Samples were vortexed for 30 seconds followed by centrifugation at 12,000 xg for 5 mins/ 4°C. Samples were then transferred using glass Pasteur pipettes into MR tubes. 1D 1H NMR spectra were acquired using a Bruker Advance III HD 700MHz spectrometer with a 5 mm TCI cryoprobe at pH 7.4/25°C. 1D 1H spectra were acquired for each sample along with water suppression, using the following Bruker pulse sequence cpmgpr1d (Carr-Purcell-Meiboom-Gill [CPMG]) with 256 transients a 15ppm spectral width, 32K points, and a 3.1 sec acquisition time. Full experimental parameters and pulse sequences are available with the deposited data at www.ebi.ac.uk/MetaboLights [2](metabolights ID MTBLS 6212)

*Spectral Processing and Data Analysis*

Initial processing of each spectrum was carried out in Bruker Topspin v3.2. Chenomx v8.2 software (www.chenomx.com, Canada) was used for metabolite annotation with identification against in house metabolite library. The spectra were normalized to the total NMR signal intensity from each sample and integrals bucketed per peak into metabolite specific intensities using in house server 'tameNMR' built within galaxy toolkit (https://github.com/PGB-LIV/tameNMR).[3] R Studio (www.r-project.org) and MetaboAnalyst 5.0 (www.metaboanalyst.ca, Canada) were used for statistical analysis. Multivariate analysis was performed, including principal component analysis (PCA) and partial least square discriminant analysis (PLS-DA). For supervised PLSDA cross validation was performed to determine model robustness on a training set of 70% of the data with area under the receiver operating curve (ROC) presented for each group with respect to all other groups in the model.[4] A threshold of >1.0 variable importance in projection (VIP) score was used to assess the discriminating metabolites. To elucidate the relevance of metabolites and metabolic pathways, MetaboAnalyst 5.0 was used for pathway enrichment analysis.[5] Once a list of metabolites was uploaded, enrichment analysis was performed using the Small Molecule Pathway Database, and then subjected to over representation analysis (ORA) using the hypergeometric test to evaluate whether a particular metabolite set is represented more than expected by chance within the given compound list. Based on the VIP loadings of PLS-DA analysis, only metabolites with a VIP >2.0 were analysed for enrichment, and one-tailed p values were used after adjusting for multiple testing.

**3D Spheroids**

*Spheroid formation*

10 µl of spheroid/mounting media was drawn up into a fluorinated ethylene propylene (FEP) tube (S 1815-04, BOLA, Germany) mounted in 50% Matrigel, 40% filtered media, 10% 25mM HEPES. For JAS239 treated spheroids, a final concentration of 500 nM JAS239 was added to the mounting media before addition to spheroids. To mimic hypoxic conditions dimethyloxalylglycine (DMOG) was added to a final concentration of 0.5 mM in mounting media. The FEP tube was inserted into a 1.5 mm glass capillary (Brand, Germany, catalogue #701908) and placed into the light-sheet sample holder, pre-set to 37°C.

*Image Acquisition and Analysis*

Spheroids were imaged on a Zeiss Light-Sheet microscope using a 561 nm laser for excitation and a 10x illumination objective. Emitted light was collected through 576–615 nm filter using a 20x W Plan-Apochromat objective. A Z-stack of 7 µm steps was acquired every 3 min for 320 cycles. Light-Sheet Z.1 Zen software was used using dual side fusion with a pivot scan. Images were detected using pco.edge scientific complementary metal–oxide–semiconductor (sCMOS) camera. Cell tracking was performed using Imaris v9.6 software (www.imaris.oxinst.com; Oxford Instruments, UK). A schematic outlining the basic pipeline of image analysis is shown in **Supplementary Figure 10**. To discount any drift of the spheroids on tracking parameters, a reference frame was applied to each frame (Supplementary Figure 2 A). Next, alignment of 'spots' (Supplementary Figure 2 B) to each cell nuclei within the spheroid were applied and quality of spots were filtered using the following parameters: average size: 10 µM, max gap of 2 at 10 µM. These parameters were then applied to the whole data set and 'dragon tail' tracks are shown (Supplementary Figure 2 C) with the colour scale denoting track duration. Data were filtered for all tracks greater than 2 h long and various parameters were plotted, including: average track speed, straightness, length and mean squared displacement (MSD).

#### References


1. Kutner, R.H., X.Y. Zhang, and J. Reiser, *Production, concentration and titration of pseudotyped HIV-1-based lentiviral vectors.* Nat Protoc, 2009. **4**(4): p. 495-505.
2. Haug, K., et al., *MetaboLights: a resource evolving in response to the needs of its scientific community.* Nucleic Acids Res, 2020. **48**(D1): p. D440-D444.
3. Al-Mutawa, Y.K., et al., *Effects of hypoxic preconditioning on neuroblastoma tumour oxygenation and metabolic signature in a chick embryo model.* Biosci Rep, 2018. **38**(4).
4. Westerhuis, J.A.H., H.C.J; Smit, S; Vis, D.J; Smilde, A.k; van Velzen, E.J.J; van Duijnhoven, J.P.M; van Dorsten, F.A; , *Assessment of PLSDA cross validation.* Metabolomics, 2008. **4**: p. 81-89.
5. Pang, Z., et al., *MetaboAnalyst 5.0: narrowing the gap between raw spectra and functional insights.* Nucleic Acids Res, 2021. **49**(W1): p. W388-W396.

#### Supplementary Figures

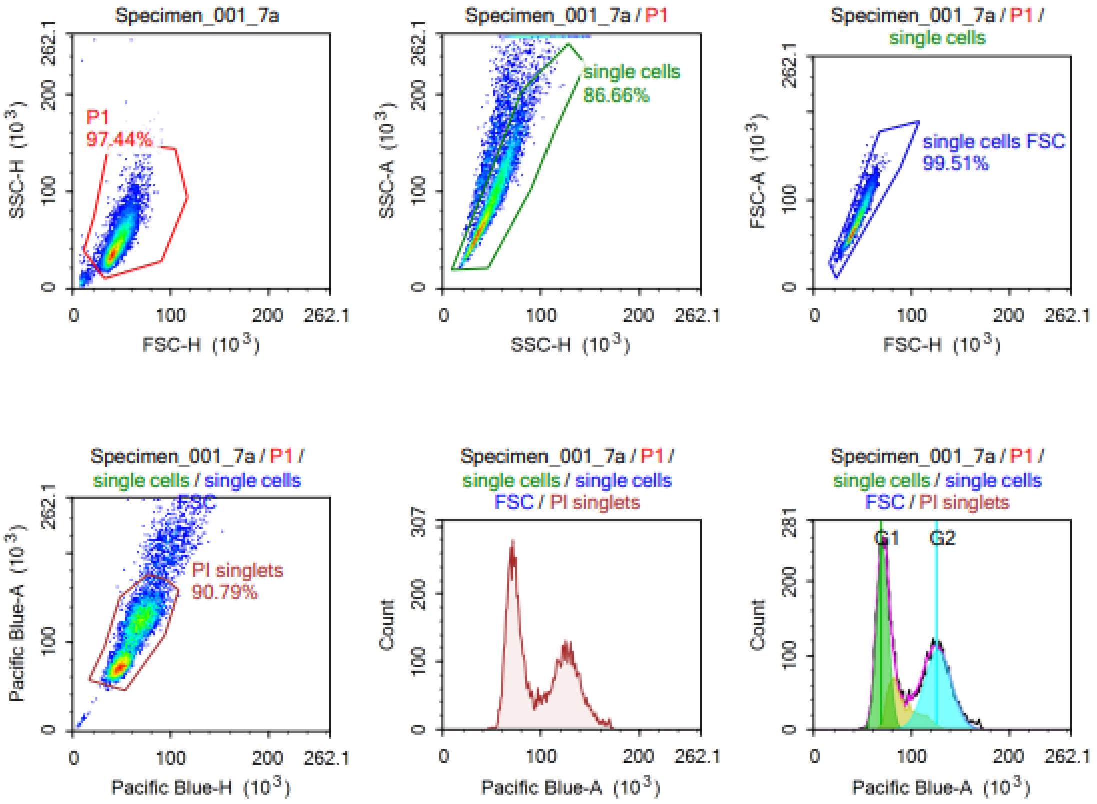


**Figure S1**. Flow cytometry data was analysed using NovoExpress 1.4.1 software. The population of cells analysed were gated to ensure only live singlets were included in the analysis and NovoExpress built-in cell cycle analysis software was used to quantify the proportion of cells in G0/G1, S and G2/M phase.

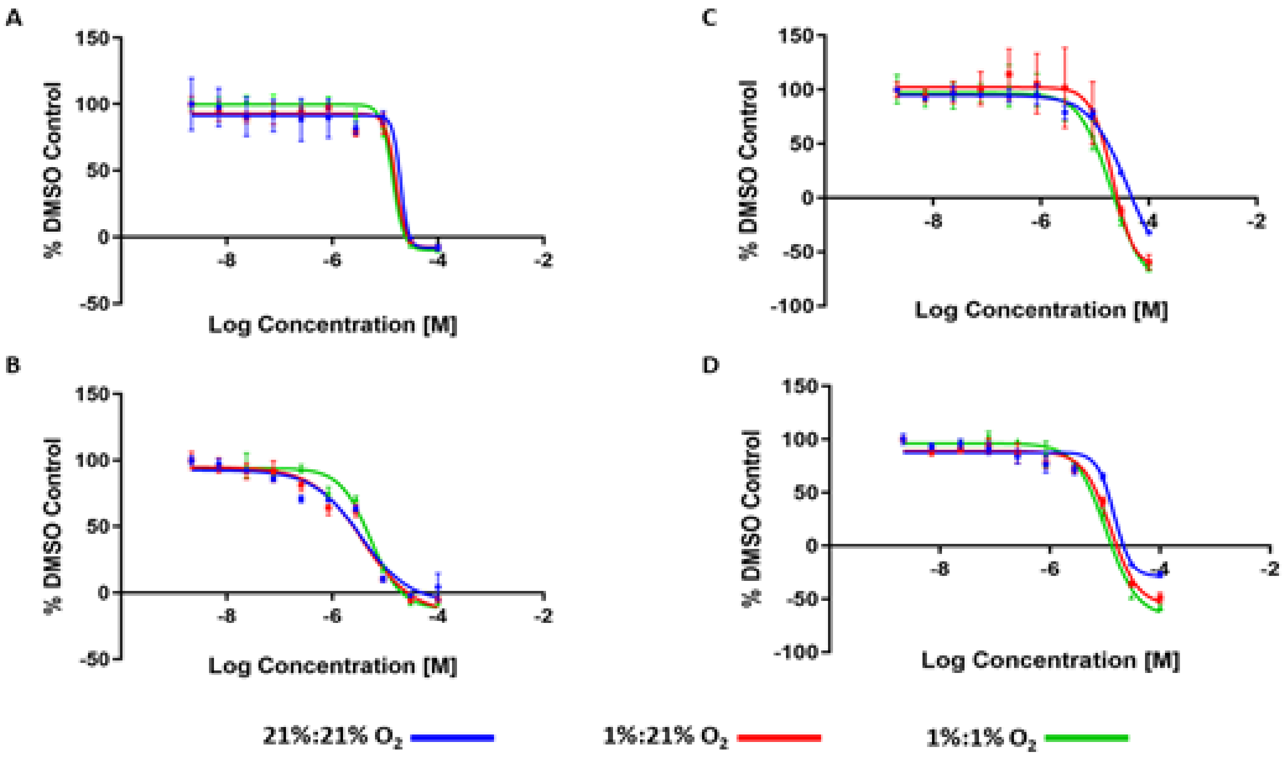


**Figure S2. MTT of F98 (A) 9L (B) U-87 MG (C) U-251 MG (D) dose response curves upon JAS239 treatment.** Half-log serial dilutions of JAS239, starting from 100µM were applied to F98, 9L, U-87 MG and U-251 MG cells that had either been preconditioned in 21% $O_2$ or 1% $O_2$ for 3 days. JAS239 was applied for 24 hours in either 21% $O_2$ (blue) or cells that were previously incubated in 1% $O_2$ for 3 days and then reoxygenated during JAS239 treatment (red) or cells preconditioned for 3 days and then treated in 1% $O_2$ for an additional 24h (green). Cellular metabolic activity was measured by MTT. Data were normalized to DMSO control; N=3.

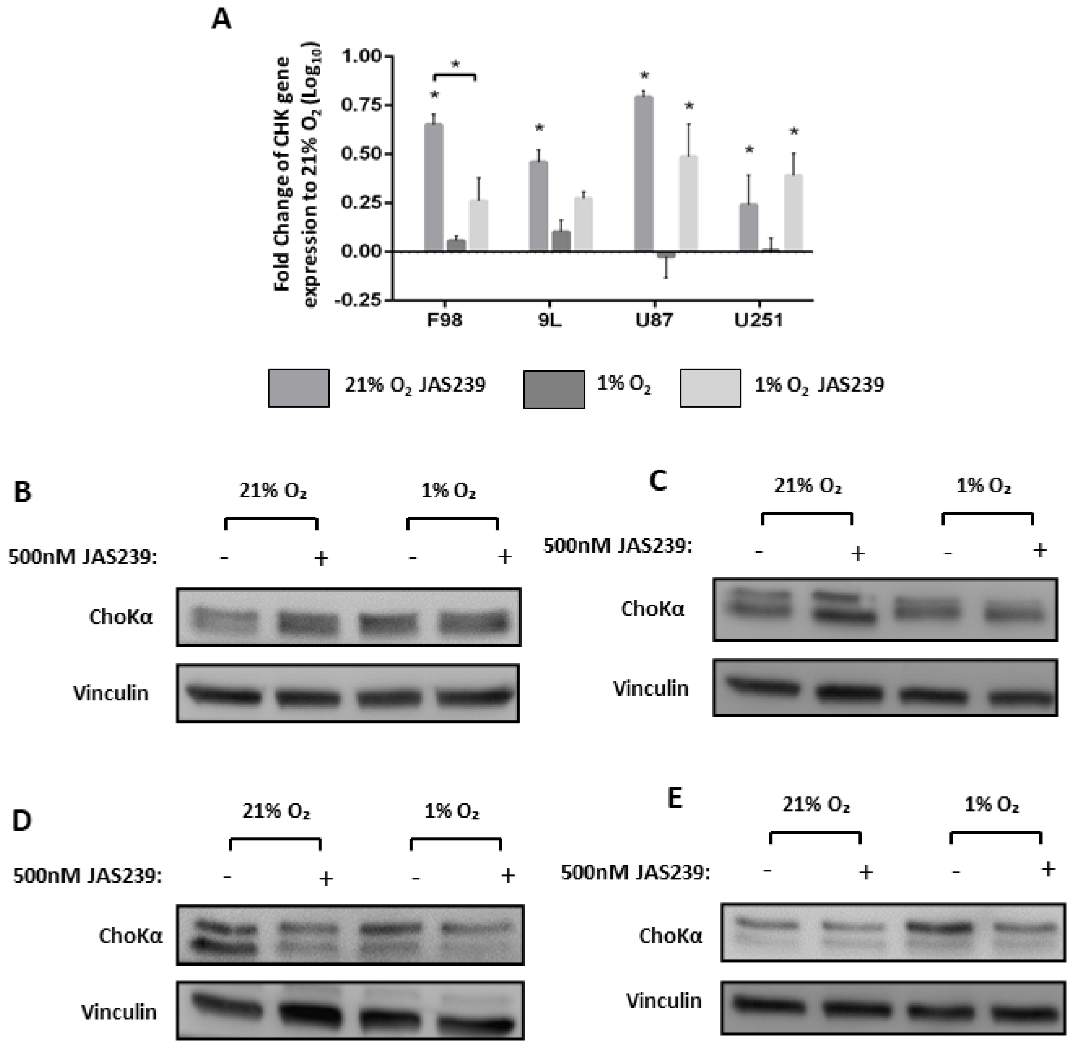


**Figure S3.** JAS239 significantly increased CHK gene expression (A) in all cell lines under normoxic conditioning (F98 $p$<0.0001, 9L $p$<0.001, U-87 MG $p$<0.0001 and U-251 MG $p$<0.05). Only U-87 MG ($p$<0.01) and U-251 MG ($p$<0.05) cells had significant increase in gene expression under hypoxic conditioning. Protein expression of F98 (B) and 9L (C) showed increased ChoKα expression with JAS239 treatment in normoxic conditioning only. U-87 MG (D) protein expression decreased with JAS239, most notably in normoxic conditioning and U-251 MG cells (E) demonstrated a reduction in hypoxic conditioning only.

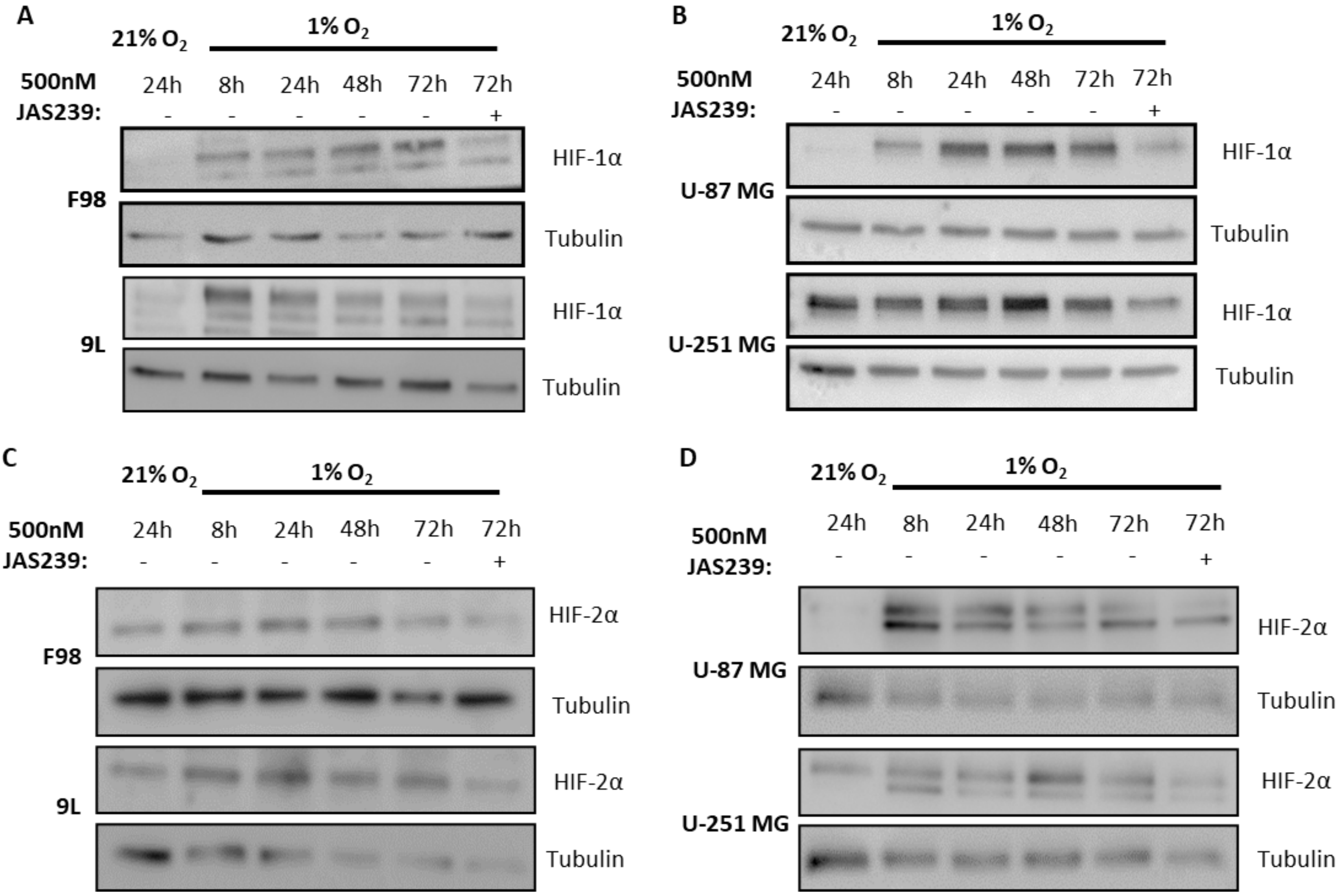


**Figure S4.** HIF-1α and HIF-2α protein expression was assessed over 72 hours and shows an induction in HIF-1α in response to hypoxia in all cell lines apart from U251 cells, where HIF-1α expression was already present in 21% $O_2$, and HIF-2α induction in all cell lines. 500nM JAS239 was added for 24 hours at the 72-hour time point. HIF-1α levels were reduced in all cell lines in presence of JAS239 (n=3).

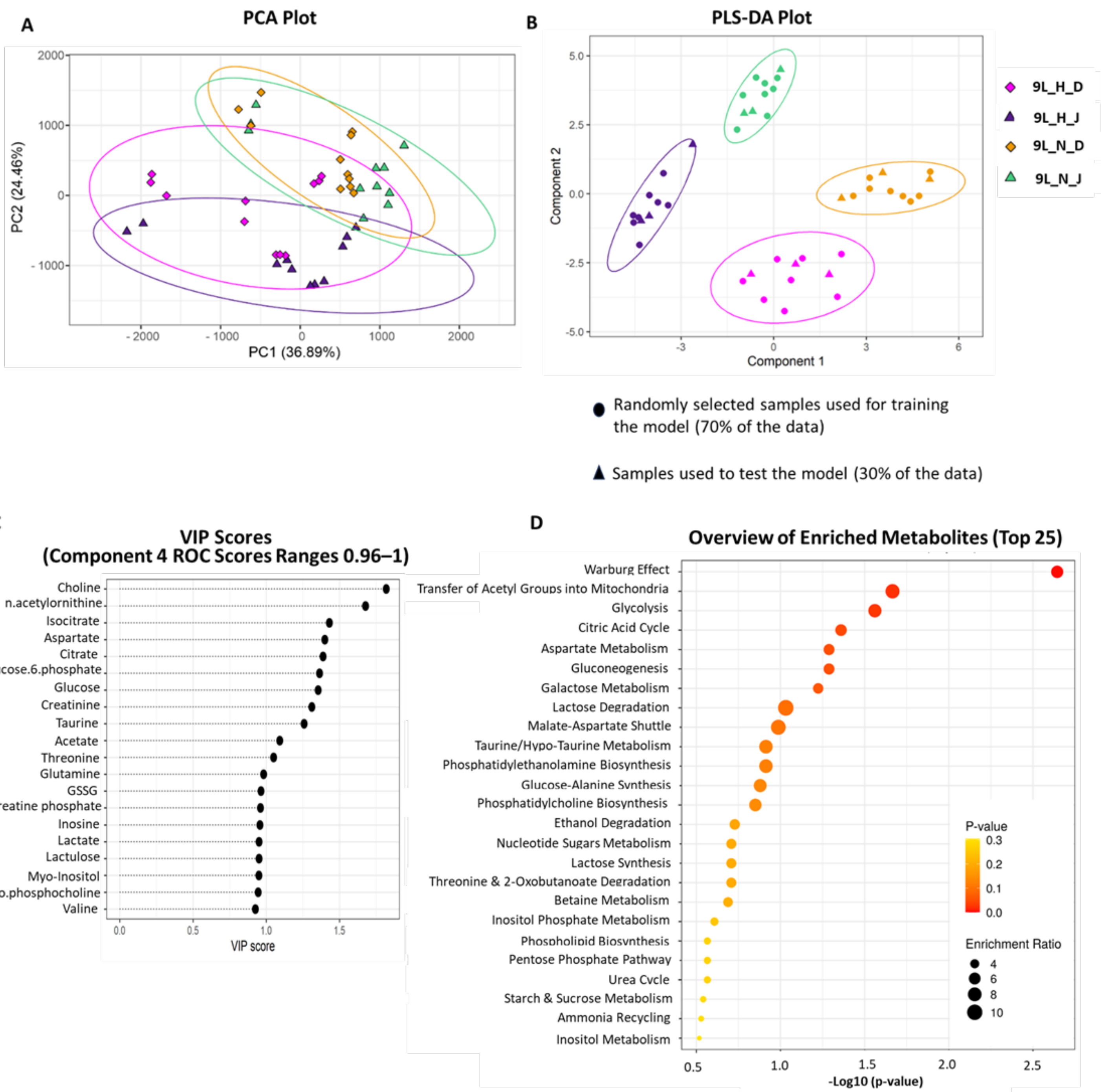


**Figure S5. 9L metabolomic analysis pipeline.** A) PCA analysis, B) PLS-DA analysis (component 4 ROC scores for each group vs all other groups: H_D = 1; H_J = 0.96; N_D = 1 and N_J = 0.96) and C) most influential metabolites. Metabolites with VIP >2 (C) informed the metabolite set enrichment analysis of top 20 metabolites using the hypergeometric test D).

D, DMSO; H, hypoxia; J; JAS239; N, normoxia; PCA, principal component analysis; PLS-DA, partial least square discriminant analysis; ROC, receiver operating characteristic; VIP, variable importance in projection coefficients.

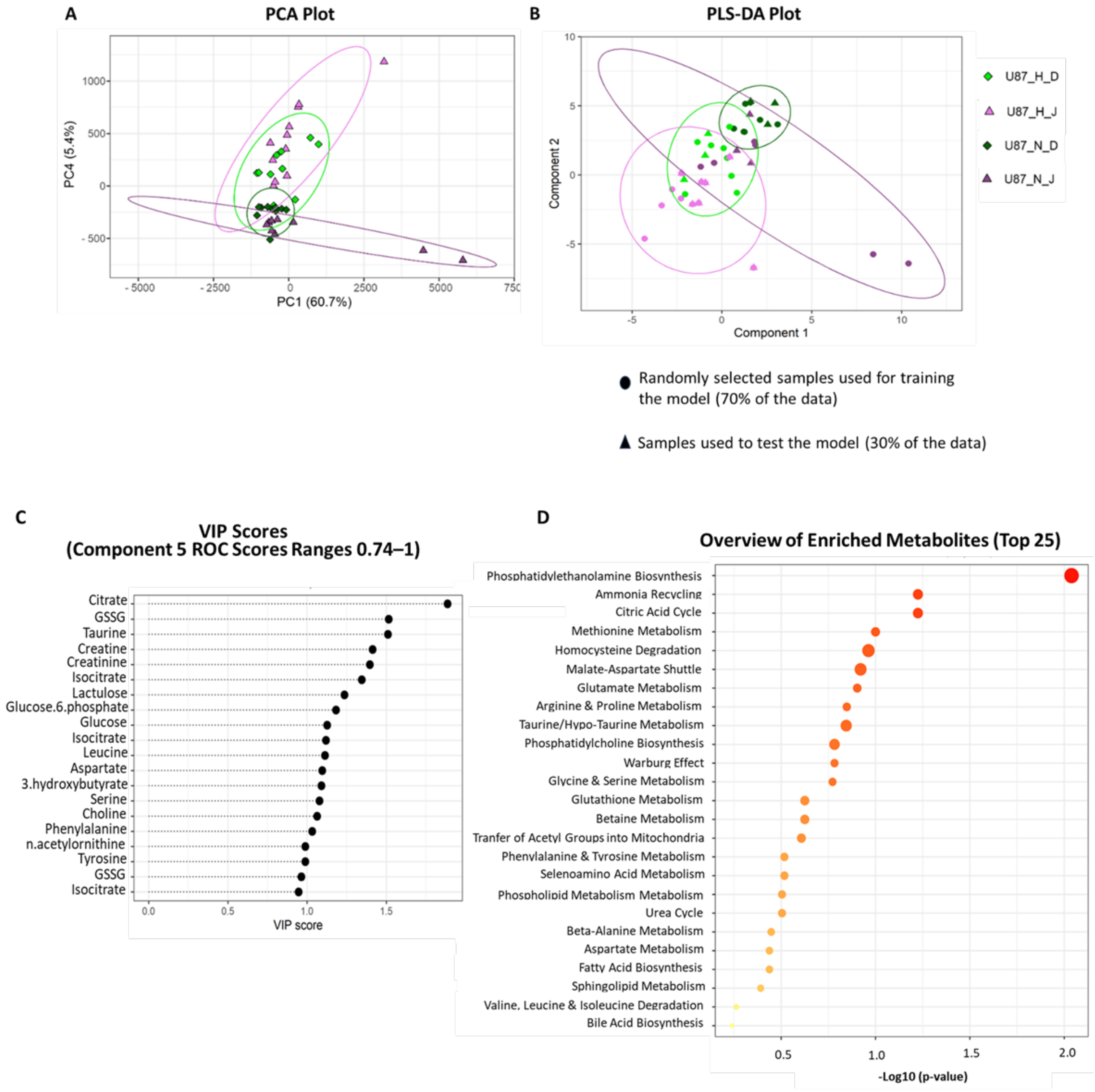


**Figure S6. U-87 MG metabolomic analysis pipeline.** A) PCA analysis, B) PLS-DA analysis (component 5 ROC scores for each group vs all other groups: H_D = 0.74; H_J = 1; N_D = 0.97 and N_J = 1) and C) most influential metabolites. Metabolites with VIP >2 (C) informed the metabolite set enrichment analysis of top 20 metabolites using the hypergeometric test D).

D, DMSO; H, hypoxia; J; JAS239; N, normoxia; PCA, principal component analysis; PLS-DA, partial least square discriminant analysis; ROC, receiver operating characteristic; VIP, variable importance in projection coefficients.

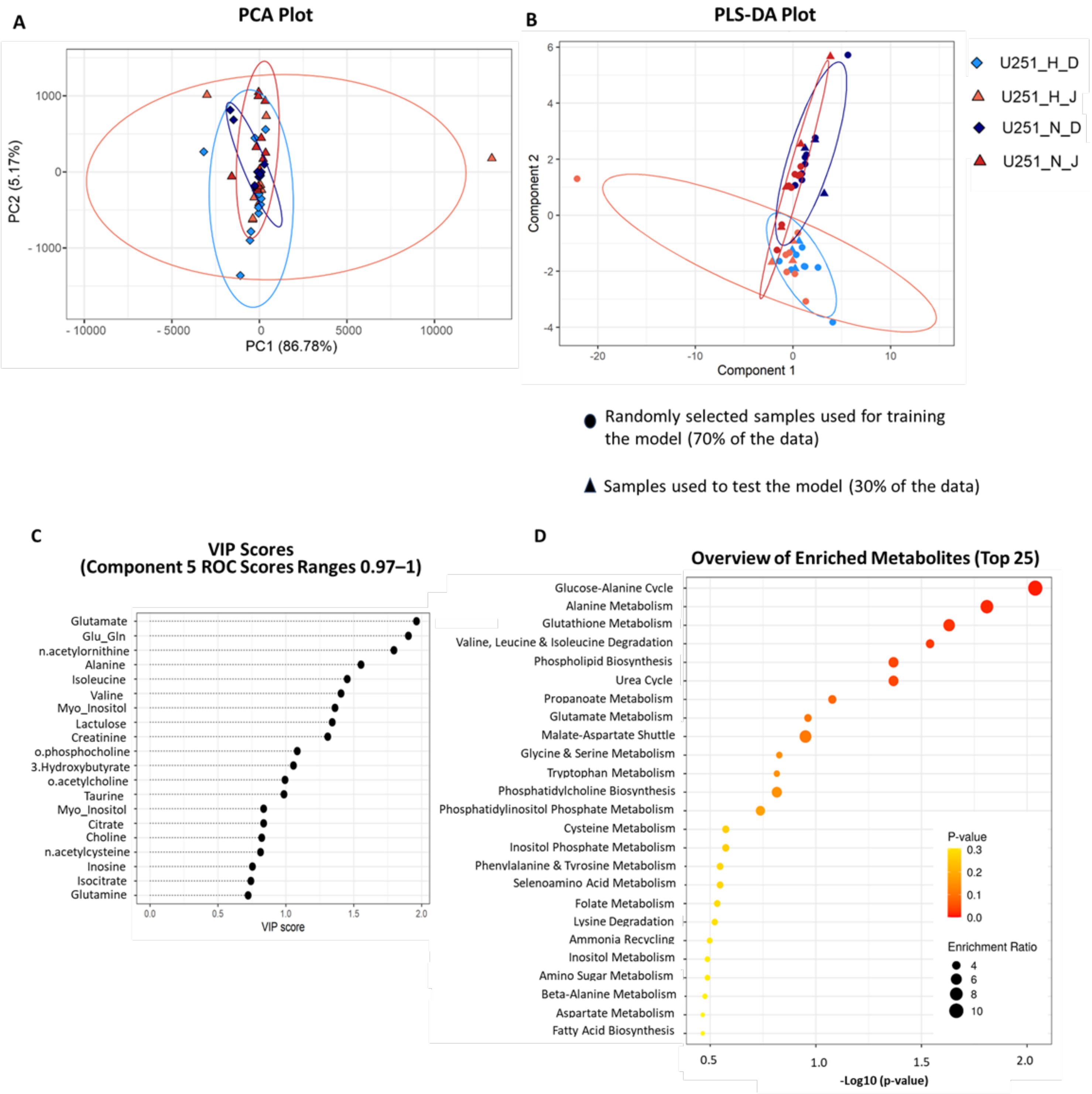


**Figure S7. U-251 MG metabolomic analysis pipeline** A) PCA analysis, B) PLS-DA analysis (component 5 ROC scores for each group vs all other groups: H_D = 0.97; H_J = 1; N_D = 1 and N_J = 1) and C) most influential metabolites. Metabolites with VIP >2 (C) informed the metabolite set enrichment analysis of top 20 metabolites using the hypergeometric test D). D, DMSO; H, hypoxia; J; JAS239; N, normoxia; PCA, principal component analysis; PLS-DA, partial least square discriminant analysis; ROC, receiver operating characteristic; VIP, variable importance in projection coefficients.

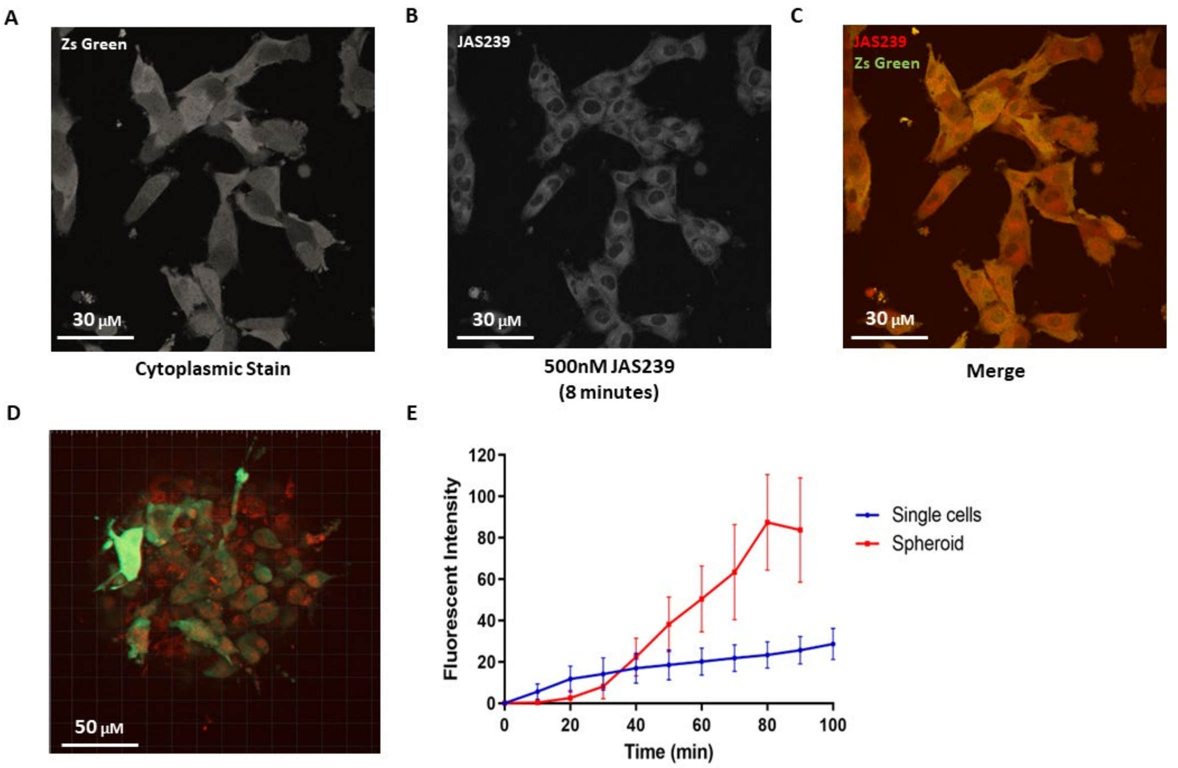


**Figure S8.** U-87 MG Zs Green labelled cells (green) were imaged on an Andor Dragonfly every 30 seconds for 2 hours in either a 2D (A–C) or 3D (D) model using 40x oil-based objective. 500nM JAS239 (red) was applied after the first z-stack was acquired. Image analysis was done using Imaris, after background subtraction, fluorescent intensity over time was plotted; FOV=3, n=3 (E).

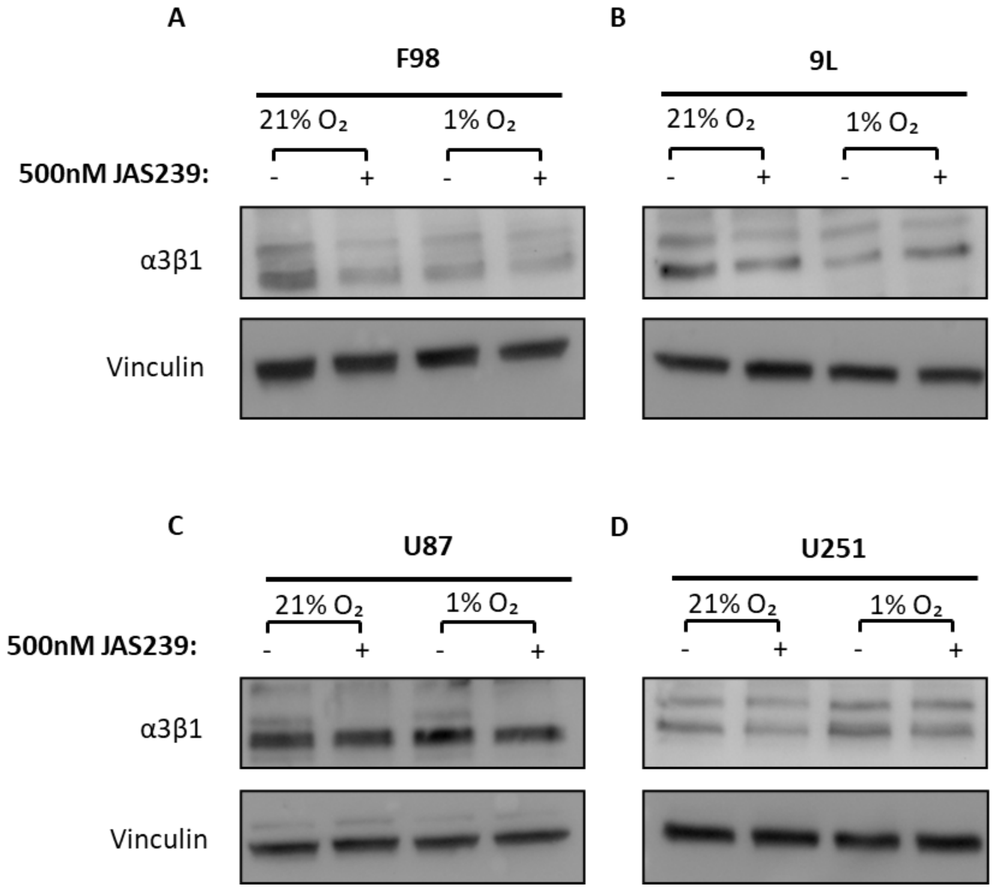


**Figure S9**. Cells were exposed to 21% or 1% $O_2$ for 96 hours before extraction. Protein expression of α3β1 for all cell lines are visible A–D.

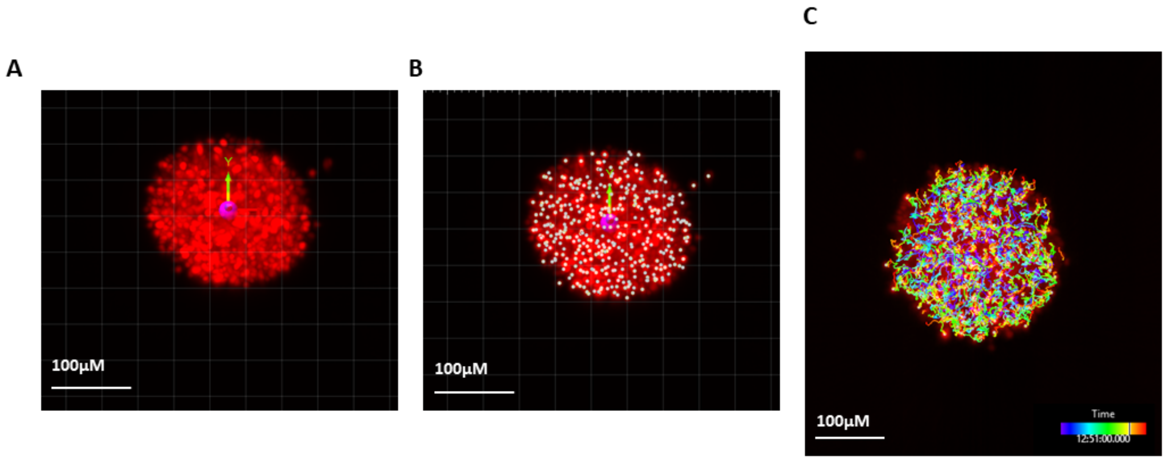


**Figure S10**. Imaris V9.6 was used to analyse cell straightness, speed, track length, and mean-squared displacement. To account for spheroid drift, a reference frame is added to all frames (A) before spots are added to the cell nuclei (B) which can be filtered based on size and gap distance between frames. Once these parameters have been applied to the whole data set, track length can be observed with ‘dragon tail’ tracks (C), the colour code inferred to the track duration.
